# DART-ID increases single-cell proteome coverage

**DOI:** 10.1101/399121

**Authors:** Albert T. Chen, Alexander Franks, Nikolai Slavov

## Abstract

Analysis by liquid chromatography and tandem mass spectrometry (LC-MS/MS) can identify and quantify thousands of proteins in microgram-level samples, such as those comprised of thousands of cells. This process, however, remains challenging for smaller samples, such as the proteomes of single mammalian cells, because reduced protein levels reduce the number of confidently sequenced peptides. To alleviate this reduction, we developed Data-driven Alignment of Retention Times for IDentification (DART-ID). DART-ID implements principled Bayesian frameworks for global retention time (RT) alignment and for incorporating RT estimates towards improved confidence estimates of peptide-spectrum-matches. When applied to bulk or to single-cell samples, DART-ID increased the number of data points by 30 – 50% at 1% FDR, and thus decreased missing data. Benchmarks indicate excellent quantification of peptides upgraded by DART-ID and support their utility for quantitative analysis, such as identifying cell types and cell-type specific proteins. The additional datapoints provided by DART-ID boost the statistical power and double the number of proteins identified as differentially abundant in monocytes and T-cells. DART-ID can be applied to diverse experimental designs and is freely available at http://github.com/SlavovLab/DART-ID.

**Author Summary:** Identifying and quantifying proteins in single cells gives researchers the ability to tackle complex biological problems that involve single cell heterogeneity, such as the treatment of solid tumors. Mass spectrometry analysis of peptides can identify their sequence from their masses and the masses of their fragment ion, but often times these pieces of evidence are insufficient for a confident peptide identification. This problem is exacerbated when analyzing lowly abundant samples such as single cells. To identify even peptides with weak mass spectra, DART-ID incorporates their retention time – the time when they elute from the liquid chromatography used to physically separate them. We present both a novel method of aligning the retention times of peptides across experiments, as well as a rigorous framework for using the estimated retention times to enhance peptide sequence identification. Incorporating the retention time as additional evidence leads to a substantial increase in the number of samples in which proteins are confidently identified and quantified.

## Introduction

Advancements in the sensitivity and discriminatory power of protein mass-spectrometry (MS) have enabled the quantitative analysis of increasingly limited amounts of samples. Recently, we have developed Single Cell Proteomics by Mass Spectrometry (SCoPE-MS). SCoPE-MS uses a barcoded carrier to boost the MS signal from single-cells and enhance sequence identification[1, 2]. While this design allows quantifying hundreds of proteins in single mammalian cells, sequence identification remains challenging because many lowly abundant peptides generate only a few fragment ions that are insufficient for confident identification [3, 4]. Such low confidence peptides are generally not used for protein quantification, and thus reduce the data points available for further analyses. We sought to overcome this challenge by using both the retention time (RT) of an ion and its MS/MS spectra to achieve more confident peptide identifications. To this end, we developed a novel data-driven Bayesian framework for aligning RTs and for updating peptide confidence. DART-ID minimizes assumptions, aligns RTs with median residual error below 3 seconds, and increases the fraction of cells in which peptides are confidently identified.

Multiple existing approaches – including Skyline ion matching [5], moFF match-between-runs [6], MaxQuant match-between-runs [7, 8], DeMix-Q [9] and Open-MS FFId [10] – allow combining MS1 spectra with other informative features, such as RT and precursor ion intensity, to enhance peptide identification. These methods, in principle, may identify any ion detected in a survey scan (MS1 level) even if it was not sent for fragmentation and second MS scan (MS2) in every run. Thus by not using MS2 spectra, these methods may overcome the limiting bottleneck of tandem MS: the need to isolate, fragment and analyze the fragments in order to identify and quantify the peptide sequence.

However not using the MS2 spectra for identification has a downside: The MS2 spectra contain highly informative features even for ions that could not be confidently identified based on spectra alone. This is particularly important when MS/MSed ions are the only ones that can be quantified, as in the case of isobaric mass tags. Thus, the MS1-based methods have a strong advantage when quantification relies only on MS1 ions (e.g., LFQ [11], and SILAC [12]), while methods using all MS2 spectra can more fully utilize all quantifiable data from isobaric tandem-mass-tag experiments.

DART-ID aims to use all MS2 spectra, including those of very low confidence PSMs, and combines them with accurate RT estimates to update peptide-spectrum-match (PSM) confidence within a principled Bayesian framework. Unlike previous MS2-based methods which incorporate RT estimates into features for FDR recalculation [13], discriminants [14], filters [15–17], or scores [18, 19], we update the ID confidence directly with a Bayesian framework [20, 21]. Crucial to this method is the accuracy of the alignment method; the higher the accuracy of RT estimates, the more informative they are for identifying the peptide sequence.

The RT of a peptide is a specific and informative feature of its sequence, and this specificity has motivated approaches aiming to estimate peptide RTs. These approaches either (i) predict RTs from peptide sequences or (ii) align empirically measured RTs. Estimated peptide RTs have a wide range of uses, such as scheduling targeted MS/MS experiments [22], building efficient inclusion and exclusion lists for LC-MS/MS [23, 24], or augmenting MS2 mass spectra to increase identification rates [14–19].

Peptide RTs can be estimated from physical properties such as sequence length, constituent amino acids, and amino acid positions, as well as chromatography conditions, such as column length, pore size, and gradient shape. These features predict the relative hydrophobicity of peptide sequences and thus RTs for LC used with MS [25–31]. The predicted RTs can be improved by implementing machine learning algorithms that incorporate confident, observed peptides as training data [15, 19, 32–35]. Predicted peptide RTs are mostly used for scheduling targeted MS/MS analyses where acquisition time is limited, e.g., multiple reaction monitoring [22]. They can also be used to aid peptide sequencing, as exemplified by “peptide fingerprinting” – a method that identifies peptides based on an ion’s RT and mass over charge (m/z) [28, 36–38]. While peptide fingerprinting has been successful for low complexity samples, where MS1 m/z and RT space is less dense, it requires carefully controlled conditions and rigorous validation with MS2 spectra [37–41]. Predicted peptide RTs have more limited use with data-dependent acquisition, i.e., shot-gun proteomics. They have been used to generate data-dependent exclusion lists that spread MS2 scans over a more diverse subset of the proteome [23, 24], as well as to aid peptide identification from MS2 spectra, either by incorporating the RT error (difference between predicted and observed RTs) into a discriminant score [14], or filtering out observations by RT error to minimize the number of false positives selected [15–17]. In addition, RT error has been directly combined with search engine scores [18, 19]. Besides automated methods of boosting identification confidence, proteomics software suites such as Skyline allow the manual comparison of measured and predicted RTs to validate peptide identifications [5].

The second group of approaches for estimating peptide RTs aligns empirically measured RTs across multiple experiments. Peptide RTs shift due to variation in sample complexity, matrix effects, column age, room temperature and humidity. Thus, estimating peptide RTs from empirical measurements requires alignment that compensates for RT variation across experiments. Usually, RT alignment methods align the RTs of two experiments at a time, and typically utilize either a shared, confidently-identified set of endogenous peptides, or a set of spiked-in calibration peptides [42, 43]. Pairwise alignment approaches must choose a particular set of RTs that all other experiments are aligned to, and the choice of that reference RT set is not obvious. Alignment methods are limited by the availability of RTs measured in relevant experimental conditions, but can result in more accurate RT estimates when such empirical measurements are available [7, 8, 43]. Generally, RT alignment methods provide more accurate estimations than RT prediction methods, discussed earlier, but also generally require more extensive data and cannot estimate RTs of peptides without empirical observations.

Methods for RT alignment are various, and range from linear shifts to non-linear distortions and time warping [44]. Some have argued for the necessity of non-linear warping functions to correct for RT deviations [45], while others have posited that most of the variation can be explained by simple linear shifts [46]. More complex methods include multiple generalized additive models [47], or machine-learning based semi-supervised alignments [48]. Once experiments are aligned, peptide RTs can be predicted by applying experiment-specific alignment functions to the RT of a peptide observed in a reference run.

Peptide RTs estimated by alignment can be used to schedule targeted MS/MS experiments – similar to the use of predicted RTs estimated from the physical properties of a peptide [43]. RT alignments are also crucial for MS1 ion/feature-matching algorithms, as discussed earlier [5–10], as well as in targeted analyses of results from data-independent acquisition (DIA) experiments [49–51]. The addition of a more complex, non-linear RT alignment model that incorporates thousands of endogenous peptides instead of a handful of spiked-in peptides increased the number of identifications in DIA experiments by up to 30% [52].

With DART-ID, we implement a novel global RT alignment method that takes full advantage of SCoPE-MS data, which feature many experiments with analogous samples run on the same nano-LC (nLC) system [1, 2]. These experimental conditions yield many RT estimates per peptide with relatively small variability across experiments. In this context, we used empirical distribution densities that obviated assumptions about the functional dependence between peptide properties, RT, and RT variability and thus maximized the statistical power of highly reproducible RTs. This approach increases the number of experiments in which a peptide is identified with high enough confidence and its quantitative information can be used for analysis.

## Results

### Model for global RT alignment and PSM confidence update

Using RT for identifying peptide sequences starts with estimating the RT for each peptide, and we aimed to maximize the accuracy of RT estimation by optimizing RT alignment. Many existing methods can only align the RTs of two experiments at a time, i.e., pairwise alignment, based on partial least squares minimization, which does not account for the measurement errors in RTs [53]. Furthermore, the selection of a reference experiment is non-trivial, and different choices can give quantitatively different alignment results. In order to address these challenges, we developed a global alignment method, sketched in Fig. 1a,b. The global alignment infers a *reference RT* for the *i*^*th*^ peptide, *μ*_*i*_ as a latent variable with value *μ*_*ik*_ in the *k*^*th*^ experiment. This can be related to the measured RT for peptide *i* in experiment *k*, *ρ*_*ik*_.

**Figure 1.**
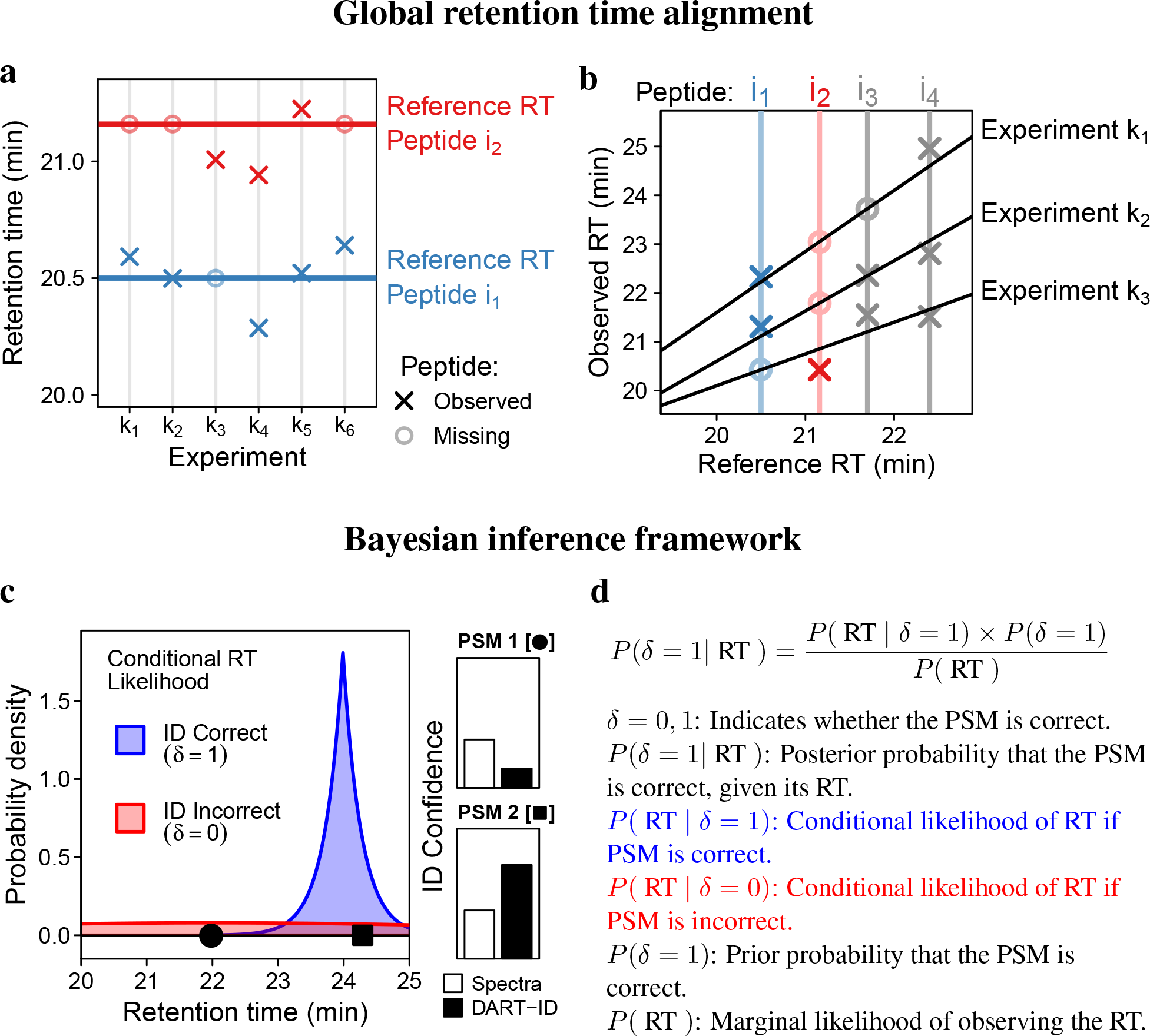
Bayesian framework for global RT alignment and matching spectra to peptides. (**a**) DART-ID defines the global reference RT as a latent variable, Eq. 1. (**b**) The observed RTs are modeled as a function of the reference RT, which allows incorporating experiment specific weights and the uncertainty in measured RTs and peptide identification as shown in Eq. 3. Then the global alignment model simultaneously infers the reference RT and aligns all experiments by solving Eq. 4. (**c**) A conceptual diagram for updating the confidence in a peptide-spectrum-match (PSM). The probability to observe each PSM is estimated from the conditional likelihoods for observing the RT if the PSM is assigned correctly (blue density) or incorrectly (red density). For PSM 1, *P* (*δ* = 1 RT) < *P* (*δ* = 0 RT), and thus the confidence decreases. Conversely, for PSM 2, *P* (*δ* = 1 RT) > *P* (*δ* = 0 RT), and thus the confidence increases. (**d**) The Bayes’ formula used to formalize the model from panel c and to update the error probability of PSMs.

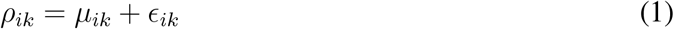

where 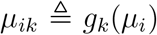 and ∊_*ik*_ is an independent mean-zero error term expressing residual (unmodeled) RT variation. As a first approximation, we assume that the observed RTs for any experiment can be well approximated using a two-segment linear regression model:

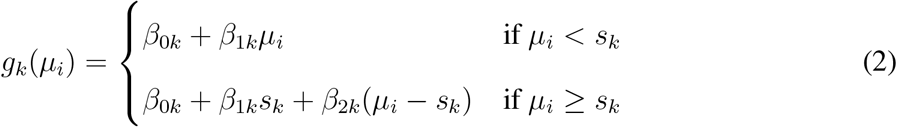

where *s*_*k*_ is the split point for the two segment regression in each experiment, and the parameters are constrained to not produce a negative RT and can be generalized to more complex monotonically-constrained models, such as spline fitting or locally estimated scatterplot smoothing (LOESS). We chose this model since we found that it outperformed a single-slope linear model by capturing more of the inter-experiment variation in RTs, Fig. S2. Based on this model, we can express the marginal likelihood for the RT of the *i*^*th*^ peptide in the *k*^*th*^ experiment as a mixture model weighted by the probability of correct sequence assignment (*λ*_*ik*_, the spectral posterior error probability (PEP)); see Fig. S1 for more details.

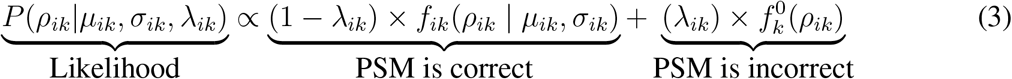

where *f*_*ik*_ is the inferred RT density for peptide *i* in experiment *k* and 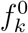 is the null RT density. In our implementation, we let 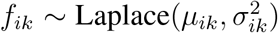 and 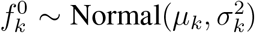 which we found worked well in practice (See Fig. S4). This framework is modular and can be easily extended to use different distributions. To account for the fact that residual RT variation increases with mean RT *and* varies between experiments (Fig. S3), we model its standard deviation, *σ*_*ik*_, as a linearly increasing function of *μ*_*i*_, Eq. 7.

Using the vectorized likelihood function from Eq. 3 and the priors described in Methods, we solve Eq. 4 to infer the joint posterior distribution of all reference RTs (and associated model parameters) across all experiments:

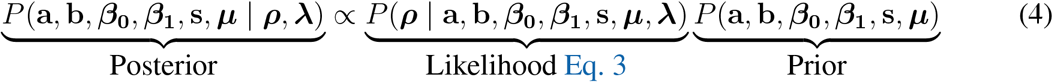

The Bayesian inference described above infers all reference RTs, ***μ***, from one global solution of Eq. 4. It allows the alignment to take advantage of any peptide observed in at least two experiments, regardless of the number of missing observations in other experiments. Furthermore, the mixture model described in Eq. 3 allows for the incorporation of low confidence peptides by using appropriate weights and accounting for the presence of false positives. Thus this method maximizes the data used for alignment and obviates the need for spiked-in standards. Furthermore, the reference RT provides a principled choice for a reference (rather than choosing a particular experiment) that is free of measurement noise. The alignment process accounts for the error in individual observations by inferring a per peptide RT distribution, as opposed to aligning to a point estimate, as well as for variable RT deviations in experiments by using experiment-specific weights.

The conceptual idea based on which we incorporate RT information for sequence identification is illustrated in Fig. 1c and formalized with Bayes’ theorem in Fig. 1d. We start with a peptide-spectrum-match (PSM) from a search engine and its associated probability to be incorrect (PEP; posterior error probability) and correct, 1-PEP. If the RT of a PSM is far from the RT of its corresponding peptide, as PSM1 in Fig. 1c, then the spectrum is more likely to be observed if the PSM is incorrect, and thus we can decrease its confidence. Conversely, if the RT of a PSM is very close to the RT of its corresponding peptide, as PSM2 in Fig. 1c, then the spectrum is more likely to be observed if the PSM is correct, and thus we can increase its confidence. To estimate whether the RT of a PSM is more likely to be observed if the PSM is correct or incorrect, we use the conditional likelihood probability densities inferred from the alignment procedure in Eq. 3 (Fig. 1b). Combining these likelihood functions with Bayes’ theorem in Fig. 1d allows to formalize this logic and update the confidence of analyzed PSMs, which we quantify with DART-ID PEPs.

### Global alignment process reduces RT deviations

To evaluate the global RT alignment by DART-ID, we used a staggered set of 46 60-minute LC-MS/MS runs performed over a span of 3 months. Each run was a diluted 1 × *M* injection of a bulk 100 × *M* SCoPE-MS sample, as described in Table 1 and by Specht et al. [2]. The experiments were run over a span of three months so that the measured RTs captured expected variance in the chromatography. The measured RTs were compared to RTs predicted from peptide sequences [30, 31, 34], and to top-performing aligning methods [7, 8, 43, 52], including the reference RTs from DART-ID; see Fig. 2a. All methods estimated RTs that explained the majority of the variance of the measured RTs, Fig. 2a. As expected, the alignment methods provided closer estimates, explaining over 99% of the variance.

**Table 1.** Design of 100 × M and 1 × M SCoPE-MS sets. Experimental design for the SCoPE-MS sets as described by Specht et al., 2018 [2], copied with permission from the authors. Schematic for the design of 100 × M sets and the proteome amounts corresponding to 1 × M sets. Jurkat and U-937 cells are cell lines of T-cells and monocytes, respectively.

| Label (TMT tag) | 100xM set | 1xM set |
| --- | --- | --- |
| 126 | 5,000 Jurkat cells | 50 Jurkat cells |
| 127N | 5,000 U-937 cells | 50 U-937 cells |
| 127C | *empty* | *empty* |
| 128N | *empty* | *empty* |
| 128C | 100 Jurkat cells | 1 Jurkat cell |
| 129N | 100 U-937 cells | 1 U-937 cell |
| 129C | 100 Jurkat cells | 1 Jurkat cell |
| 130N | 100 U-937 cells | 1 U-937 cell |
| 130C | 100 Jurkat cells | 1 Jurkat cell |
| 131N | 100 U-937 cells | 1 U-937 cell |
| 131C | *empty* | *empty* |

**Figure 2.**
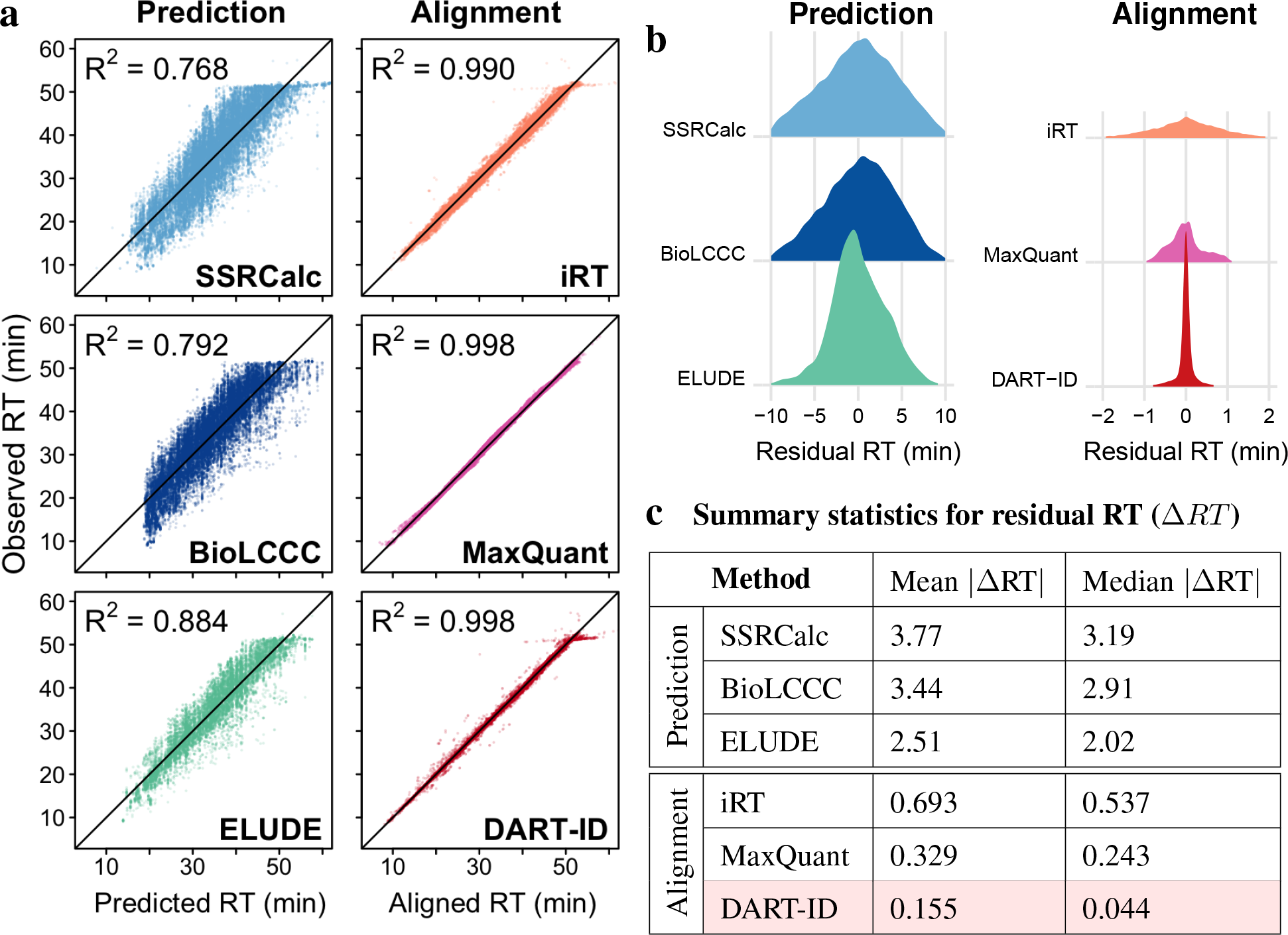
Comparison of inferred reference RTs to empirical RTs. (**a**) Scatter plots of observed RTs versus inferred RTs. The comparisons include 33,383 PSMs with PEP < 0.01 from 46 LC-MS/MS runs over the span of three months. The left column displays comparisons for RT prediction methods – SSRCalc [30], BioLCCC [31], and ELUDE [34]. The right column displays comparisons for alignment methods – precision iRT [52], MaxQuant match-between-runs [7, 8], and DART-ID. (**b**) Distributions of residual RTs: ∆*RT* = Observed RT − Reference RT. Note the different scales of the x-axes between the prediction and alignment methods. (**c**) Mean and median of the absolute values of ∆*RT* from panel (b).

To evaluate the accuracy of RT estimates more rigorously, we compared the distributions of differences between the reference RTs and measured RTs, shown in Fig. 2b. This comparison again underscores that the differences are significantly smaller for alignment methods, and smallest for DART-ID. We further quantified these differences by computing the mean and median absolute RT deviations, i.e., |∆RT|, which is defined as the absolute value of the difference between the observed RT and the reference RT. For the prediction methods – SSRCalc [30], BioLCCC [31], and ELUDE [34] – the average deviations exceed 2 min, and ELUDE has the smallest average deviation of 2.5 min. The alignment methods result in smaller average deviations, all below < 1 min, and DART-ID shows the smallest average deviation of 0.044 min (2.6 seconds).

### DART-ID increases proteome coverage of SCoPE-MS experiments

Search engines such as MaxQuant [7, 8] use the similarity between theoretically predicted and experimentally measured MS2 spectra of ions to match them to peptide sequences, i.e., peptide-spectrum-matches (PSM). The confidence of a PSM is commonly quantified by the probability of an incorrect match: the posterior error probability (PEP) [21, 54, 55]. Since the estimation of PEP does not include RT information, we sought to update the PEP for each PSM by incorporating RT information within the Bayesian framework displayed in Fig. 1c,d. This approach allowed us to use the estimated RT distributions for each peptide with minimal assumptions.

The Bayesian framework outlined in Fig. 1c,d can be used with RTs estimated by other methods, and its ability to upgrade PSMs is directly proportional to the accuracy of the estimated RTs. To explore this possibility, we used our Bayesian model with RTs estimated by all methods shown in Fig. 2. The updated error probabilities of PSMs indicate that all RT estimates enhance PSM discrimination, Fig. S5. Even lower accuracy RTs predicted from peptide sequence can be productively used to upgrade PSMs. However, the degree to which PSMs are upgraded, i.e. the magnitude of the confidence shift, increases with the accuracy of the RT estimates and is highest with the DART-ID reference RTs.

We refer to the PEP assigned by the search engine (MaxQuant throughout this paper) as “Spectral PEP”, and after it is updated by the Bayesian model from Fig. 1d as “DART-ID PEP”. Comparing the Spectral and DART-ID PEPs indicates that the confidence for some PSMs increases while for others decreases; see density plot in Fig. 3a. Reassuringly, all PSMs with low Spectral PEPs have even lower DART-ID PEPs, meaning that all confident PSMs become even more confident. On the other extreme, many PSMs with high Spectral PEPs have even higher DART-ID PEPs, meaning that some low-confidence PSMs are further downgraded. Confidence upgrades, where DART-ID PEP < Spectral PEP, range within 1–3 orders of magnitude.

**Figure 3.**
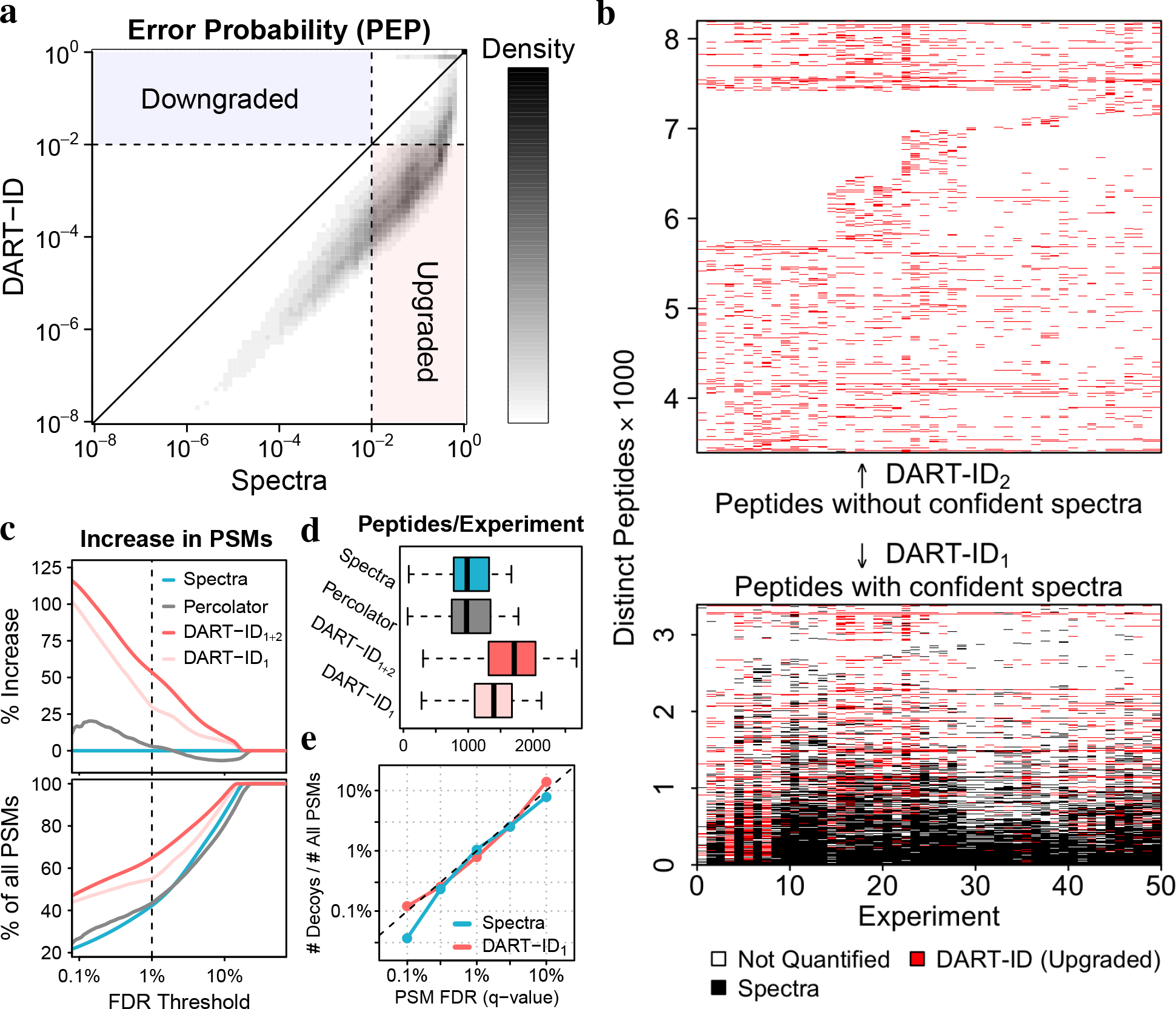
Incorporating RTs increases confident peptide identifications. (**a**) A 2D density distribution of error probabilities derived from spectra alone (Spectral PEP), compared to that after incorporating RT evidence (DART-ID PEP). (**b**) Map of all peptides observed across all experiments. Black marks indicate peptides with Spectral FDR < 1%, and red marks peptides with DART-ID FDR < 1%. (**c**) Increase in confident PSMs (top), and in the fraction of all PSMs (bottom) across the confidence range of the x-axis. The curves correspond to PEPs estimated from spectra alone, from spectra and RTs using percolator and from spectra and RTs using DART-ID. DART-ID identifications are split into DART-ID_1_ and DART-ID_2_ depending on whether the peptides have confident spectral PSMs as marked in panel (b). (**d**) Distributions of number of unique peptides identified per experiment. (**e**) The fraction of decoys, i.e. the number of decoy hits divided by the total number of PSMs, as a function of the FDR estimated from spectra alone or from DART-ID. The Spectral FDR is estimated from separate MaxQuant searches, with the FDR applied on the peptide level.

The density plot in Fig. 3a displays a subset of peptides with Spectral PEP > 0.01 and DART-ID PEP < 0.01. These peptides have low confidence of identification based in their MS/MS spectra alone, but high confidence when RT evidence is added to the spectral evidence. To visualize how these peptides are distributed across experiments, we marked them with red dashes in Fig. 3b. The results indicate that the data sparsity decreases; thus DART-ID helps mitigate the missing data problem of shotgun proteomics. Fig. 3b is separated into two subsets, DART-ID1 and DART-ID2, which correspond respectively to peptides that have at least one confident spectral PSM, and peptides whose spectral PSMs are all below the set confidence threshold of 1% FDR. While the PSMs of DART-ID2 very likely represent the same peptide sequence – since by definition they share the same RT, MS1 m/z and MS2 fragments consistent with its sequence – we cannot be confident in the exact sequence assignment. Thus, they are labeled separately and their sequence assignment further validated in the next section. The majority of PSMs whose confidence is increased by DART-ID have multiple confident Spectral PSMs, and thus reliable sequence assignment. Analysis of newly identified peptides in Fig. 3c shows that DART-ID helps identify about 50% more PSMs compared to spectra alone at an FDR threshold of 1%. This corresponds to an increase of ~30 – 50% in the fraction of PSMs passing an FDR threshold of 1%, as shown in the bottom panel of Fig. 3c. Furthermore, the number of distinct peptides identified per experiment increases from an average of ~1000 to an average of ~1600, Fig. 3d. Percolator, a widely used FDR recalculation method that also incorporates peptide RTs [13], also increases identification rates, albeit to a lesser degree than DART-ID, Fig. 3c,d.

We observe that DART-ID PEPs are bimodally distributed (Fig. S6), suggesting that DART-ID acts as an efficient binary classifier. Modifying error probabilities, however, does risk changing the overall false discovery rate (FDR) of the PSM set. To evaluate the effect of DART-ID on the overall FDR, we allowed the inclusion of decoy hits in both the alignment and confidence update process [54]. The results from this analysis in Fig. 3e indicate that, as expected, the fraction of PSMs matched to decoys is proportional to the FDR estimated both from the Spectral PEP and from the updated DART-ID PEP. We encourage users of DART-ID to evaluate the results from applying DART-ID and other related methods on their datasets using this benchmark as well as the numerous quantitative benchmarks described in the subsequent sections.

### DART-ID increases proteome coverage of bulk LC-MS/MS experiments

While we were motivated to develop DART-ID within the context of the SCoPE-MS method, we show in Fig. 4 that DART-ID is similarly able to increase quantitative coverage in a label-free [56] and a TMT-labelled [57] bulk LC-MS/MS experiment. The DART-ID alignment performed differently between the label-free set (120 min gradients) and the TMT-labelled set (180 min gra-dients) Fig. 4a, with slightly higher residuals for the longer gradient. The percent increase in confident PSMs, when using DART-ID PEPs instead of spectral PEPs Fig. 4b, also fell into the expected range of 30–50% at 1% FDR. The increase in confident PSMs is shown in discrete terms in Fig. 4c, where experiments in both the label-free and TMT-labelled sets receive thousands of more confident PSMs that can then be used for further quantitative analysis.

**Figure 4.**
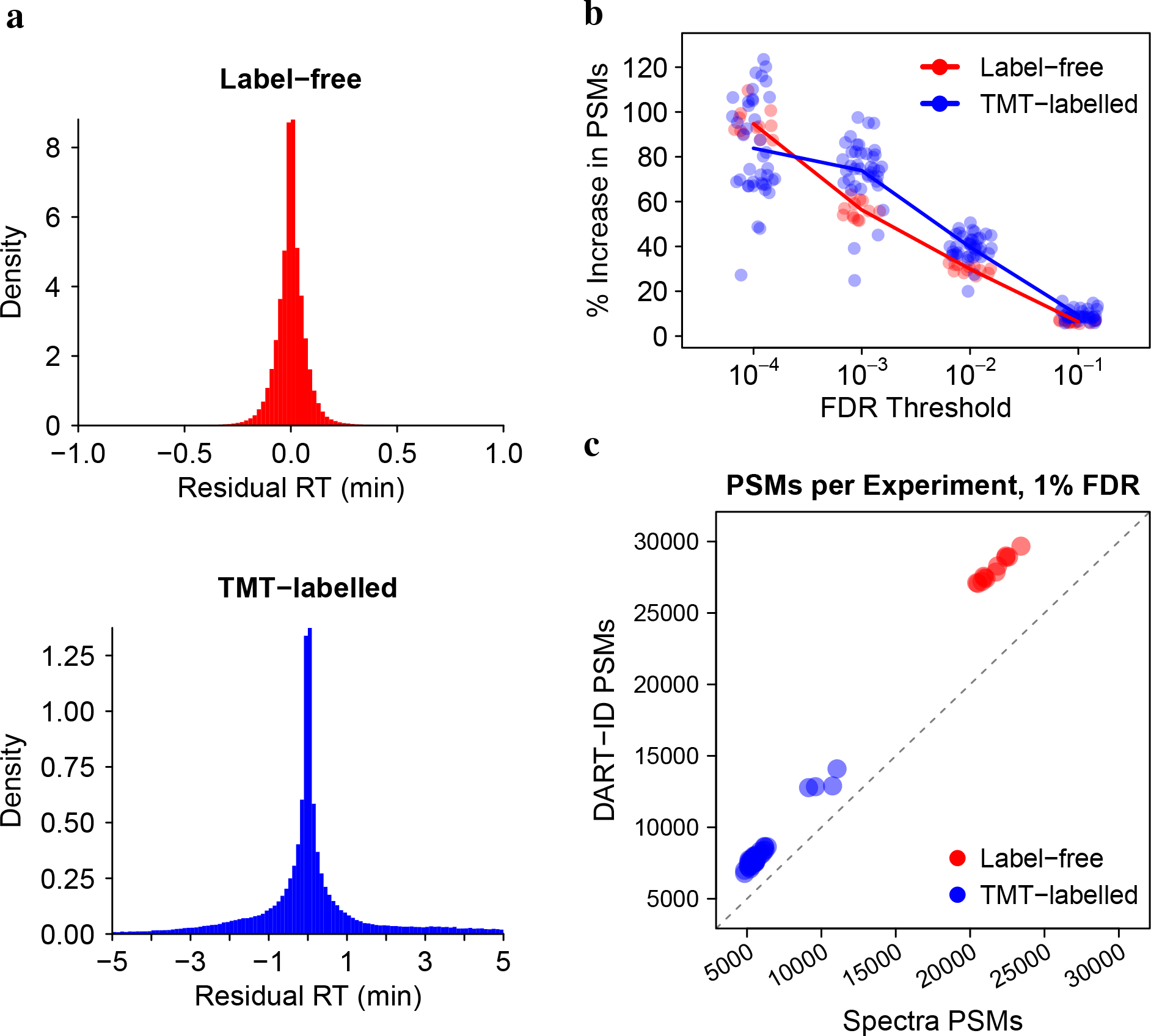
Application of DART-ID on bulk LC-MS/MS runs. Residual RTs after DART-ID alignment for (**a**) label-free dataset [56] and TMT-labelled dataset [57]. (**b**) DART-ID doubles the PSMs at 0.01% FDR and increase them by about 40% at 1% FDR. Each circle corresponds to the number of PSMs in an LC-MS/MS run. (**c**) Number of PSMs per run at 1% FDR, after applying DART-ID versus before its application. The x-coordinate represents the *Spectra* PSMs and and y-coordinate represents the *DART-ID* PSMs at 1% FDR.

### DART-ID decrease missing datapoints

These increases of confident PSMs, in both the SCoPE-MS and bulk LC-MS/MS sets, decreases the amount of missing data per run. In Fig. 5a we show qualitatively that DART-ID can fill in many of these missing values on the protein level. On the level of experimental runs, as shown quantitatively in Fig. 5b, DART-ID significantly reduces the amount of missing data and mitigates the stochasticity that is inherently to data-dependent MS methods.

**Figure 5.**
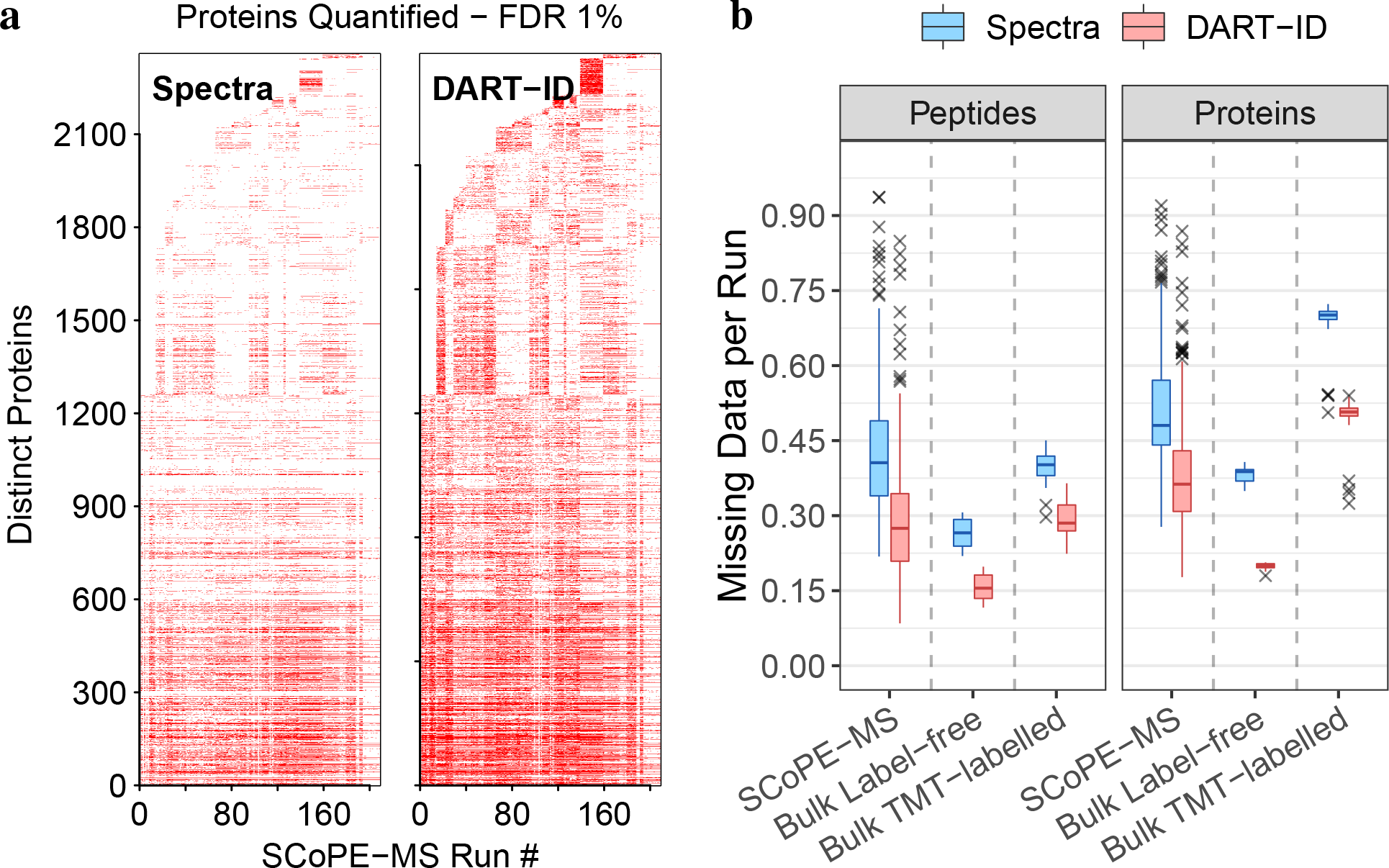
DART-ID decreases missing datapoints across runs. (**a**) Map of quantified proteins across 209 SCoPE-MS runs, before and after applying DART-ID. A red mark denotes a protein quantified in an run at 1% FDR. Only peptides seen in >50% of experiments are included. (**b**) Decrease in missing data across all runs after applying DART-ID, for SCoPE-MS and the two bulk sets from Fig. 4 at 1% FDR. All corresponding *Spectra* and *DART-ID* distributions differ significantly; the probability that they are sampled from the same distribution ≪ 1 ∗ 10^−10^.

### Validation of PSMs upgraded by DART-ID

We next sought to evaluate whether the confident DART-ID PSMs without confident Spectral PSMs, i.e. DART-ID_2_ from Fig. 3b, are matched to the correct peptide sequences. To this end, we sought to evaluate whether the RTs of such PSMs match the RTs for the corresponding peptides identified from high-quality, confident spectra. For this analysis, we split a set of experiments into two subsets, *A* and *B*, Fig. 6a. The application of DART-ID to *A* resulted in two disjoint subsets of PSMs: *A*_1_, corresponding to PSMs with confident spectra (Spectral PEP < 0.01), and *A*_2_, corresponding to “upgraded” PSMs (Spectral PEP > 0.01 and DART-ID PEP < 0.01). We overlapped these subsets with PSMs from *B* having Spectral PEP < 0.01, so that the RTs of PSMs from *B* can be compared to the RTs of PSMs from subsets *A*_1_ and *A*_2_, Fig. 6a. This comparison shows excellent agreement of the RTs for both subsets *A*_1_ and *A*_2_ with the RTs for high quality spectral PSMs from *B*, Fig. 6b,c. This result suggests that even peptides upgraded without confident spectral PSMs are matched to the correct peptide sequences.

**Figure 6.**
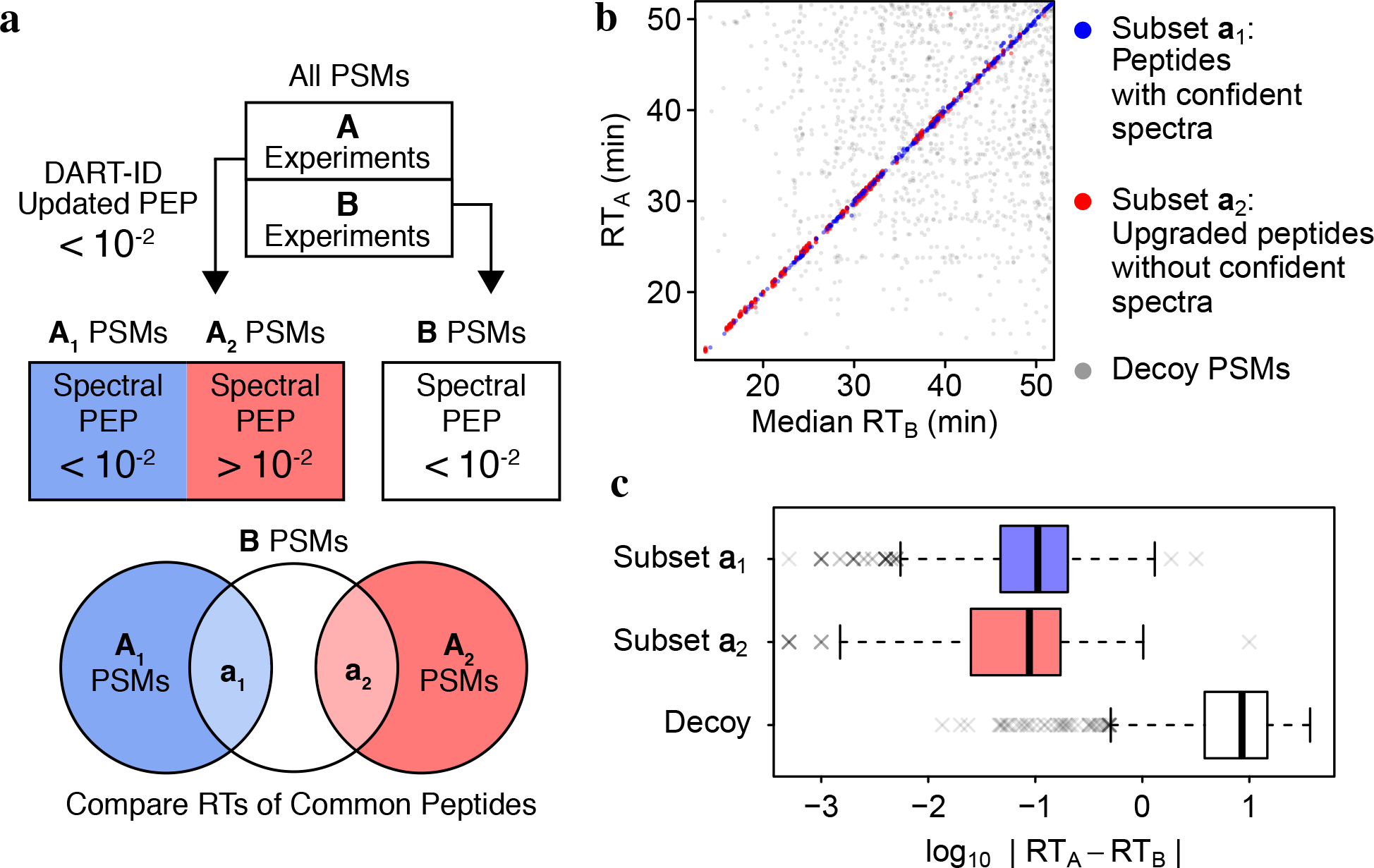
Validation of newly identified peptides with RT of technical replicates. (**a**) Schematic design of this validation experiment. It used 11 technical replicate LC-MS/MS experiments that were run on the same day. (**b**) Comparison of the RTs of subsets *a*_1_ and *a*_2_ to the RTs of corresponding peptides from *B*. Decoy PSMs have randomly sampled RTs and are included here as a null model. (**c**) Residual RT distributions for the two subsets of data *a*_1_ and *a*_2_ as defined in panel a and for a decoy subset.

### Validation by internal consistency

We ran DART-ID on SCoPE-MS method development experiments [2], all of which contain quantification data in the form of 11-plex tandem-mass-tag (TMT) reporter ion (RI) intensities. Out of the 10 TMT “channels”, six represent the relative levels of a peptide in simulated single cells, i.e., small bulk cell lysate diluted to a single cell-level level. These six single cell channels are made of T-cells (Jurkat cell line) and monocytes (U-937 cell line). We then used the normalized TMT RI intensities to validate upgraded PSMs by analyzing the consistency of protein quantification from distinct peptides.

Internal consistency is defined by the expectation that the relative intensities of PSMs reflect the relative levels of their corresponding proteins. If upgraded PSMs are consistent with Spectral PSMs for the same protein, then their relative RI intensities will have lower coefficients of variation (CV) within a protein than across different proteins [58]. CV is defined as *σ*/*μ*, where *σ* is the standard deviation and *μ* is the mean of the normalized RI intensities of PSMs belonging to the same protein. A negative control is constructed by creating a decoy dataset where PSM protein assignments are randomized.

For this and later analyses, we filter PSMs from our data into the following disjoint sets:

- *Spectra* – Spectral PEP < 0.01
- *DART-ID* – (Spectral PEP > 0.01) ∩ (DART-ID PEP < 0.01)
- *Percolator* [13] – (Spectral PEP > 0.01) ∩ (Percolator PEP < 0.01)

where *Spectra* is disjoint from the other two sets, i.e., Spectra ∩ DART-ID = ∅ and Spectra ∩ Percolator = ∅. These sets of PSMs, as depicted in Fig. 7a, are intersected with each other through a set of shared proteins between the three sets of PSMs.

**Figure 7.**
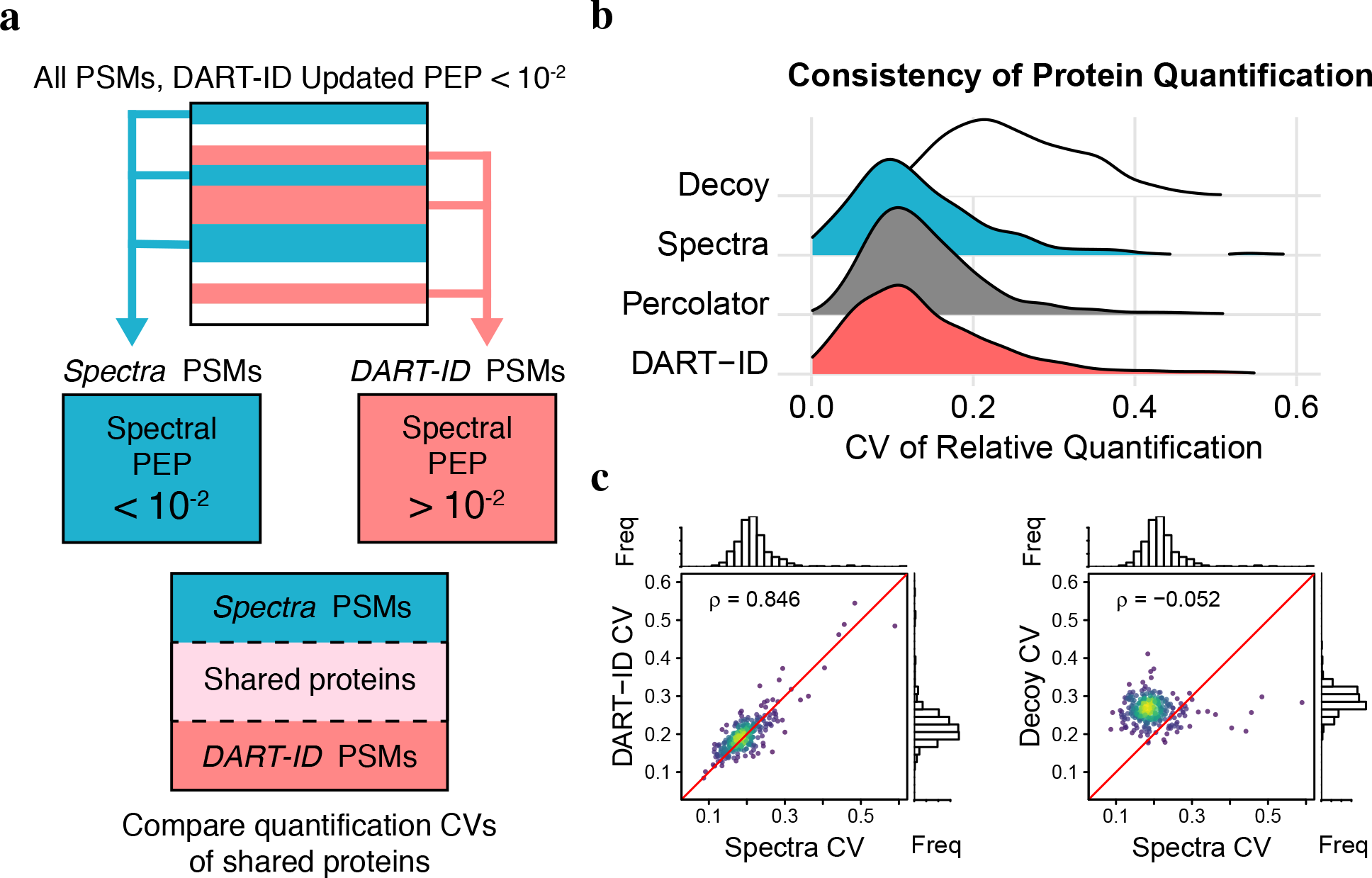
Validation of boosted PSMs by internal consistency. (**a**) Schematic for separating PSM subsets, where *Spectra* and *DART-ID* subsets of PSMs are disjoint. (**b**) Distributions of coefficient of variation (CVs) for each protein in each subset. *Decoy* is a subset of PSMs with their protein assignments randomized. (**c**) Comparing protein CVs of *n* = 275 proteins between the *Spectra* and *DART-ID* PSM subsets, and from the *Spectra* and *Decoy* subsets.

The protein CVs of the *Spectra*, *DART-ID*, and *Percolator* PSM sets, depicted in Fig. 7b, show similar distributions and smaller CVs than those from the decoy set. In addition, Fig. 7c shows agreement between the protein CVs of the *Spectra* and *DART-ID* PSM sets, as opposed to the CVs of the *Spectra* set and *Decoy* set. This demonstrates that the protein-specific variance in the relative quantification, due to either technical or biological noise, is preserved in these upgraded PSMs.

### Proteins identified by DART-ID separate cell types

The upgraded PSMs from the *DART-ID* set are not just representative of proteins already quan-tified from confident spectral PSMs, but when filtering at a given confidence threshold (e.g., 1% FDR), they allow for the inclusion of new proteins for analysis. As the quantification of these new proteins from the *DART-ID* PSMs cannot be directly compared to that of the proteins from the *Spectra* PSMs, we instead compare how the new proteins from *DART-ID* can explain the biological differences between two cell types – T-cells (Jurkat cell line) and monocytes (U-937 cell line) – present in each sample and experiment. The data was split into sets in the same manner as the previous section, as shown in Fig. 7a, where the *Spectra* and *DART-ID* sets of PSMs are disjoint. We then filtered out all PSMs from *DART-ID* that belonged to any protein represented in *Spectra*, so that the sets of proteins between the two sets of PSMs were disjoint as well.

To test whether or not DART-ID identified peptides consistently across experiments, we used principal component analysis (PCA) to separate the T-cells and monocytes quantified in our experiments. This PCA analysis in Fig. 8a shows clear separation of T-cells and monocytes from both the *Spectra* and *DART-ID* PSM sets. If boosted peptide identifications were spurious and inconsistent, then the PCA analysis could not separate the cell types or cluster them together. In addition, relative protein ratios (T-cells/monocytes) estimated from the two disjoint PSM sets are in good agreement (*ρ* = 0.84); see Fig. S7.

**Figure 8.**
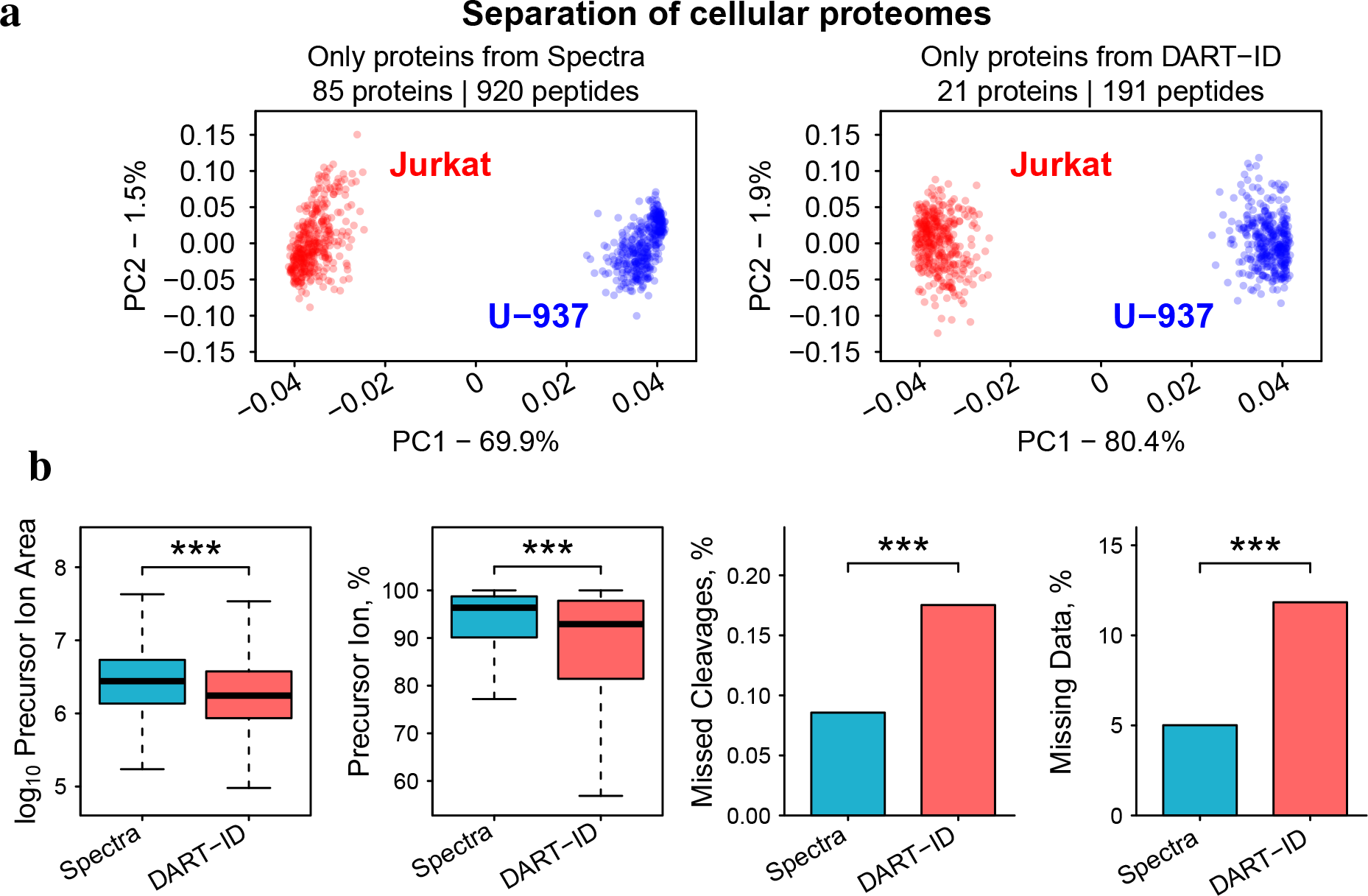
Quantification of proteins identified by spectra alone and by DART-ID. (**a**) Principal component analysis of the proteomes of 375 samples corresponding to either T-cells (Jurkat cell line) or to monocytes (U-937 cell line). The *Spectra* set contains proteins with Spectral PSMs filtered at 1% FDR, and the *DART-ID* set contains a disjoint set of proteins quantified from PSMs with high Spectral PEP but low DART-ID PEP. Only peptides with less than 5% missing data were used for this analysis, and the missing data were imputed. (**b**) The distributions of some features of the *Spectra* and *DART-ID* PSMs differ slightly. These features include: precursor ion area is the area under the MS1 elution peak and reflects peptide abundance; precursor ion fraction which reflects MS2 spectral purity; missed cleavages is the average number of internal lysine and arginine residues; and % missing data is the average fraction of missing TMT reporter ion quantitation per PSM. All distributions are significantly different, with *p* < 10^−4^.

While DART-ID_2_ PSMs are able to uncover entirely new proteins carrying consistent biological signal, on average these PSMs differ slightly from Spectral PSMs in purity, missed-cleavages, and missing data; see Fig. 8b. However, the distributions of these features are largely overlapping, and the magnitude of these differences are relatively small; most spectra of DART-ID PSMs are still >90% pure, and have less than 16% missing data and missed cleavages. Of course the intended usage of DART-ID is not to separate these two groups of PSMs and analyze them separately, but instead to combine them and increase the number of data points available for analysis. Indeed, adding DART-ID PSMs to the *Spectra* PSMs doubles the number of differentially abundant proteins between T-cells and monocytes, Fig. 9a,b,c.

**Figure 9.**
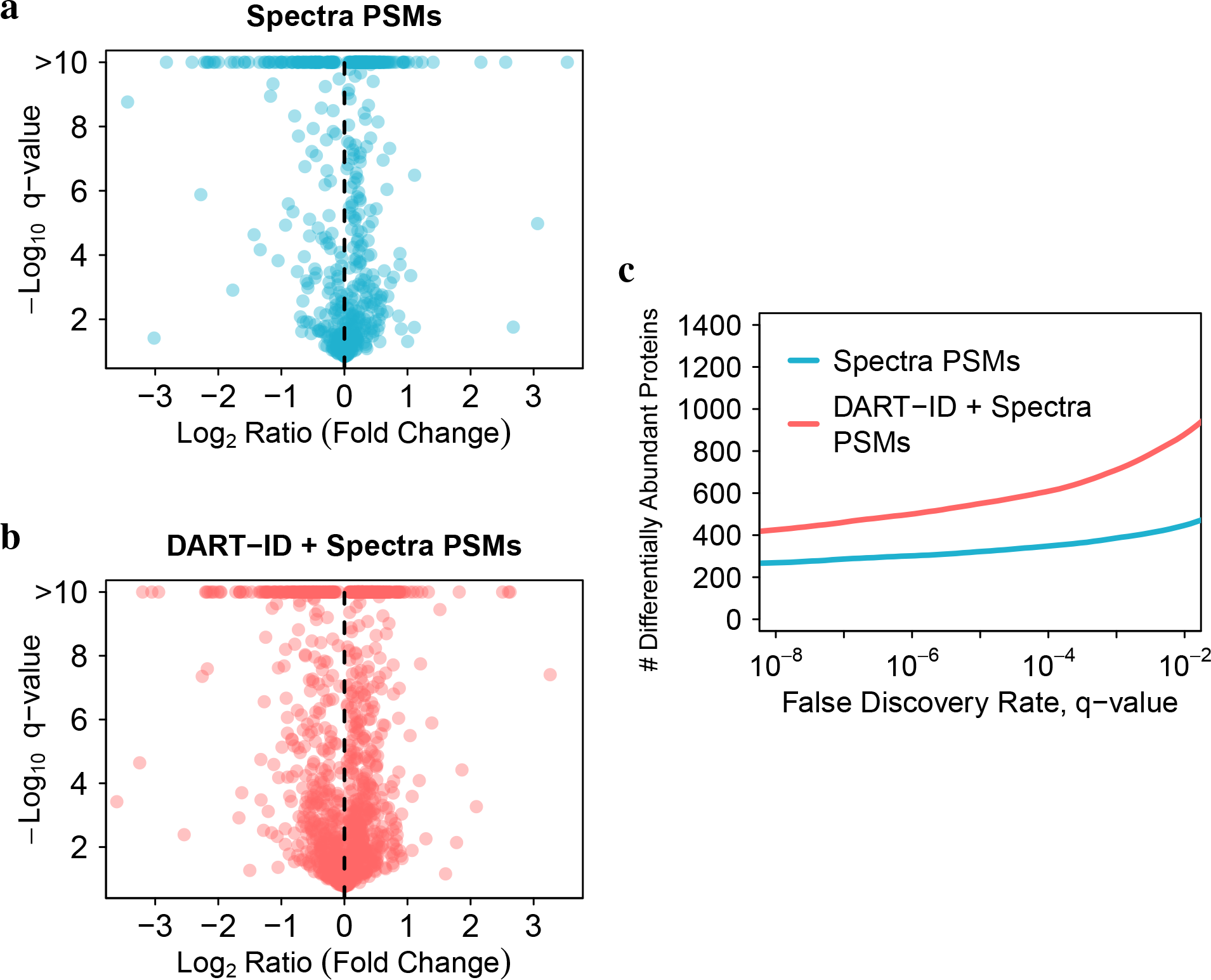
DART-ID identifies more differentially abundant proteins. The difference in protein abundance between T-cells and monocytes was visualized in the space of fold-change and its significance, i.e., volcano plots. The volcano plot using only proteins quantified from *Spectra* PSMs (**a**) identifies fewer proteins than the volcano plot using proteins from *Spectra* + *DART-ID* PSMs (**b**). Fold changes are averaged normalized RI intensities of T-cells (Jurkat cell line) / monocytes (U-937 cell line). q-values are computed from two-tailed t-test p-values and corrected for multiple hypotheses testing. (**c**) Number of differentially abundant proteins as a function of the significance FDR from panels a and b.

## Discussion

Here we present DART-ID, a new Bayesian approach that infers RTs with high accuracy and uses these accurate RT estimates to improve peptide sequence identification. We demonstrate that DART-ID can estimate and align RTs with accuracy of a few seconds for 60 minute LC-MS/MS runs and can leverage this high accuracy towards increasing the confidence in correct PSMs and decreasing the confidence in incorrect PSMs. This principled and rigorous estimation of the confidence of PSMs increases quantification coverage by 30 − 50%, primarily by increasing the number of experiments in which a peptide is quantified.

We validated the upgraded PSMs using methods for FDR estimation (Fig. 3e), cross-validation (Fig. 6), intra-protein CV validation (Fig. 7), and biological signal validation (Fig. 8). All of these methods strongly support the reliability of DART-ID inferences. We encourage the use of these methods for benchmarking the application of DART-ID (and any other related method) on other datasets.

DART-ID is applicable to any large set of LC-MS/MS analyses with consistent LC setup. The more consistent the LC, the more powerful DART-ID is since its statistical power is proportional to the accuracy of RT estimates. Our SCoPE-MS runs provide an example of runs with consistent LC [1, 4] and motivated us to develop DART-ID. However, we found (show in Fig. 4) that DART-ID performs similarly well with bulk LC-MS/MS runs of TMT-labeled and label-free samples.

A principal advantage of DART-ID is that its probabilistic model naturally adapts to the RT reproducibility and obviates thresholds, e.g., a threshold on RT errors. Rather DART-ID updates the confidence of each PSMs using a rigorous quantitative model based on empirically derived distributions of RT reproducibility. Thus, it adapts and controls for the reproducibility of the LC and the accuracy of the RT estimates as shown in Fig. S5.

Another principal advantage of DART-ID is its ability to use all PSMs (including those with sparse observations and low confidence) to create a global RT alignment. This is possible because DART-ID alignment takes into account the confidence of PSMs as part of the mixture model in Eq. 3. This results in accurate RT estimates (Fig. 2) that are robust to missing data and benefit from all PSMs regardless of their identification confidence.

If the LC and RTs of a dataset are very variable, one may extend the alignment model beyond Eq. 2 to capture the increased variability. The two-segment linear regression from Eq. 2 demonstrated here captures more variation than a single-slope linear regression. DART-ID, however, is not constrained to these two functions and can implement any monotone function. Non-linear functions that are monotonically constrained, such as the *logit* function, have been implemented in our model during development. More complex models, for example monotonically-constrained general additive models, could increase alignment accuracy further given that the input data motivates added complexity.

While DART-ID is focused on aligning and utilizing RTs from LC-MS/MS experiments, the alignment method could potentially be applied to other separation methods, including ion mobility, gas chromatography, supercritical fluid chromatography, and capillary electrophoresis. The ion drift time obtained from instruments with an ion mobility cell are particularly straightforward to align and incorporate by DART-ID’s Bayesian frameowrk. Another potential extension of DART-ID is to offline separations prior to analysis, i.e., fractionation. RT alignment would only be applicable between replicates of analogous fractions, but a more complex model could also take into account membership of a peptide to a fraction as an additional piece of evidence.

DART-ID is modular, and the RT alignment module and PEP update modules may be used separately. For example, the RT estimates may be applied to increase the performance of other peptide identification methods incorporating RT evidence 14–17. One application is integrating the inferred RT from DART-ID into the search engine score, as done by previous methods [18, 19], to change the best hit for a spectrum, save a spectrum from filtering due to high score similarities (i.e., low delta score) [21], or provide evidence for hybrid spectra. Although DART-ID’s alignment is based on point estimates of RT, the global alignment methodology could also be applied to feature-based alignments [6, 8–10] to obviate the limitations inherent in pairwise alignments.

## Methods

### Data sources and experimental design

The data used for the development and validation of the DART-ID method were 263 method-development experiments for SCoPE-MS and its related projects. All samples were lysates of the Jurkat (T-cell), U-937 (monocyte), or HEK-293 (human embryonic kidney) cell lines. Samples were prepared with the mPOP sample preparation protocol, and then digested with trypsin [2]. All experiments used either 10 or 11-plex TMT for quantification. Most but not all sets followed the experimental design as described by Table 1. All experiments were run on a Thermo Fisher (Waltham, MA) Easy-nLC system with a Waters (Milford, MA) 25cm × 75μm, 1.7μ BEH column with 130Å pore diameter, and analyzed on a Q-Exactive (Thermo Fisher) mass spectrometer. Gradients were run at 100 nL/min from 5-35%B in 48 minutes with a 12 minute wash step to 100%B. Solvent composition was 0% acetonitrile for A and 80% acetonitrile for B, with 0.1% formic acid in both. A subset of later experiments included the use of a trapping column, which extended the total run-time to 70 minutes. Detailed experimental designs and mass spectrometer parameters of each run can be found in Table S1. All Thermo .RAW files are publicly available online. More details on sample preparation and analysis methods can be found from the mPOP protocol [2].

### Searching raw MS data

Searching was done with MaxQuant v1.6.1.0 [7] against a UniProt protein sequence database with 443722 entries. The database contained only SwissProt entries and was downloaded on 5/1/2018. Searching was also done on a contaminant database provided by MaxQuant, which contained common laboratory contaminants and keratins. MaxQuant was run with Trypsin specificity which allowed for two missed cleavages, and methionine oxidation (+15.99492 Da) and protein N-terminal acetylation (+42.01056 Da) as variable modifications. No fixed modifications apart from TMT were specified. TMT was searched using the “Reporter ion MS2” quantification setting on MaxQuant, which searches for the TMT addition on lysine and the n-terminus with a 0.003 Da tolerance. Observations were selected at a false discovery rate (FDR) of 100% at both the protein and PSM level to obtain as many spectrum matches as possible, regardless of their match confidence. All raw MS files, MaxQuant search parameters, the sequence database, and search outputs are publicly available online.

### Data filtering

Only a subset of the input data is used for the alignment of experiments and the inference of RT distributions for peptides. First, decoys and contaminants are filtered out of the set. Contaminants may be problematic for RT alignment since their retention may be poorly defined, e.g., they may be poorly chromatographically resolved. Then, observations are selected at a threshold of PEP < 0.5.

Observations are additionally filtered through a threshold of retention length, which is defined by MaxQuant as the range of time between the first matched scan of the peptide and the last matched scan. Any peptide with retention length > 1 min for a 60 min run is deemed to have too wide of an elution peak, or chromatography behavior more consistent with contaminants than retention on column. In our implementation, this retention length threshold can be set as a static number or as a fraction of the total run-time, i.e., (1/60) of the gradient length.

For our data, only peptide sequences present in 3 or more experiments were allowed to participate in the alignment process. The model can allow peptides only present in one experiment to be included in the alignment, but the inclusion of this data adds no additional information to the alignment and only serves to slow it down computationally. The definition of a peptide sequence in these cases is dynamic, and can include modifications, charge states, or any other feature that would affect the retention of an isoform of that peptide. For our data, we used the peptide sequence with modifications but did not append the charge state.

Preliminary alignments revealed certain experiments where chromatography was extremely abnormal, or where peptide identifications were too sparse to enable an effective alignment. These experiments were manually removed from the alignment procedure after a preliminary run of DART-ID. From the original 263 experiments, 37 had all of their PSMs pruned, leaving only 226 experiments containing PSMs with updated confidences. These experiments are included in the DART-ID output but do not receive any updated error probabilities as they did not participate in the RT alignment. All filtering parameters are publicly available as part of the configuration file that was used to generate the data used in this paper.

### Global alignment model

Let *ρ*_*ik*_ be the RT assigned to peptide *i* in experiment *k*. In order to infer peptide and experiment-specific RT distributions, we assume that there exists a set of reference retention times, *μ*_*i*_, for all peptides *i*. Each peptide has a unique reference RT, independent of experiment. We posit that for each experiment, there is a simple monotone increasing function, *g*_*k*_, that maps the reference RT to the predicted RT for peptide *i* in experiment *k*. An observed RT can then be expressed as

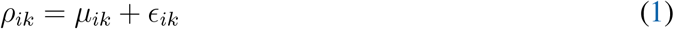

where 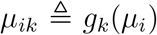 and ∊_*ik*_ is an independent mean-zero error term expressing residual (unmodeled) RT variation. As a first approximation, we assume that the observed RTs for any experiment can be well approximated using a two-segment linear regression model:

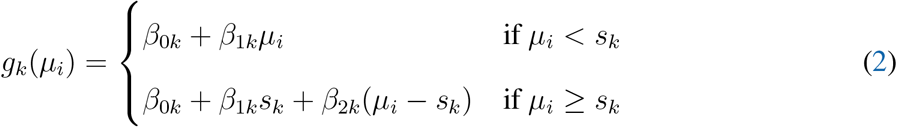

where *s*_*k*_ is the split point for the two segment regression in each experiment, and the parameters are constrained to not produce a negative RT. This two-piece model was found to outperform a single-slope linear model, Fig. S2. This model can be extended to more complex monotonic models, such as spline fitting, or non-linear monotonic models, such as a logit function or LOESS.

To factor in the spectral PEP given by the search engine, and to allow for the inclusion of low probability PSMs, the marginal likelihood of an RT in the alignment process can be described using a mixture model as described in Fig. S1. For a PSM assigned to peptide *i* in experiment *k* the RT density is

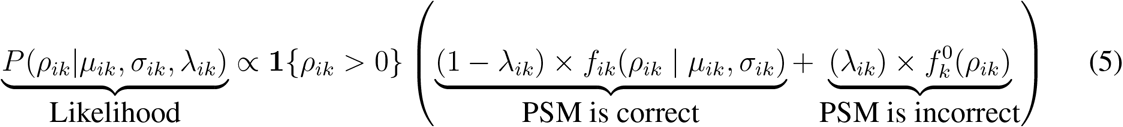

where *λ*_*ik*_ is the error probability (PEP) for the PSM returned by MaxQuant, *f*_*ik*_ is the inferred RT density for peptide *i* in experiment *k* and 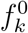 is the null RT density. In our implementation, we let:

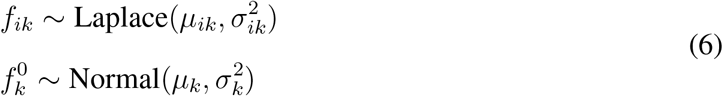

which we found worked well in practice (See Fig. S4). However, our framework is modular and it is straightforward to utilize different residual RT and null distributions if appropriate. For example, with non-linear gradients that generate a more uniform distribution of peptides across the LC run [22], it may be sensible for the null distribution to be defined as uniformly distributed, i.e. 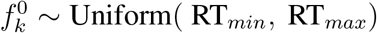.

Finally, to reflect the fact that residual RT variation increases with mean RT *and* varies between experiments (Fig. S3), we model the standard deviation of a peptide RT distribution, *σ*_*ik*_, as a linear function of the reference RT:

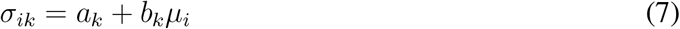

where *μ*_*i*_ is the reference RT of the peptide sequence, and *a*_*k*_ and *b*_*k*_ are the intercept and slope which we infer for each experiment. *a*_*k*_, *b*_*k*_ and *μ*_*i*_ are constrained to be positive, and hence *σ*_*ik*_ > 0 as well.

To estimate all unknown parameters, we consider the joint posterior distribution of the experiment specific alignment parameters and the reference RTs given the observed retention times,

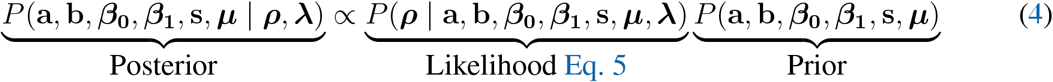

where *P*(a, b, *β*_0_, *β*_1_, s, ***μ***) are the prior distributions for all unknown alignment parameters and reference RTs and *P*(***ρ*** | a, b, *β*_0_, *β*_1_, s,***μ***) is the likelihood, as determined by Equation. a, b, *β*_0_, *β*_1_, s are all *K*-vectors of alignment parameters for each experiment. ***μ*** consists of the reference RTs for every peptide.

The priors for the bayesian inference can be found in the `.stan` model files, and for the analyses in this paper, are as follows:

Reference RT: *μ* ~ Normal(RT_mean_, RT_sd_)
Global Sigma Slope: *b*_global_ ~ Lognormal(0.1, 0.5)
Sigma Slope: *b* ~ Lognormal(log(*b*_global_), 1)
Sigma Intercept: *a* ~ Lognormal(0, 2)
Intercept: *β*_0_ ~ Normal(0, 10)
First Segment: *β*_1_ ~ Lognormal(0, 0.5)
Second Segment: *β*_2_ ~ Lognormal(0, 0.5)
Split Point: *s* ~ Uniform(0, max(RT))

where RTmean and RT_sd_ are the mean and standard deviation of all RTs across all experiments, respectively. max(RT) is the maximum observed RT of all RTs across all experiments. These priors were chosen for groups of 60 min LC-MS/MS runs, and can be adjusted accordingly for different run lengths, gradient shapes, and groupings of runs with different run times.

### Alignment Comparison

We compared the DART-ID alignment accuracy against five other RT prediction or alignment algorithms. As some methods returned absolute predicted RTs (such as BioLCCC [31]) and others returned relative hydrophobicity indices (such as SSRCalc [30]), a linear regression was built for each prediction method. Alignment accuracy was evaluated using three metrics: *R*^2^, the Pearson correlation squared, and the mean and median of |ΔRT|, the absolute value of the residual RT, and is defined as |Observed RT − Predicted RT|. We selected only confident PSMs (PEP < 0.01) for this analysis, and used data that consisted of 33383 PSMs from 46 LC-MS/MS experiments run over the course of 90 days in order to produce more chromatographic variation. A list of these experiments is found in Table S1.

SSRCalc [30] was run from SSRCalc Online (http://hs2.proteome.ca/SSRCalc/SSRCalcQ.html), with the “100Å C18 column, 0.1% Formic Acid 2015” model, “TMT” modification, and “Free Cysteine” selected. No observed RTs were inputted along with the sequences.

BioLCCC [31] was run online fromhttp://www.theorchromo.ru/ with the parameters of 250mm column length, 0.075mm column inner diameter, 130Å packing material pore size, 5% initial concentration of component B, 35% final concentration of component B, 48 min gradient time, 0 min delay time, 0.0001 ml/min flow rate, 0% acetonitrile concentration in component A, 80% ace-tontrile concentration in component B, “RP/ACN+FA” solid/mobile phase combination, and no cysteine carboxyaminomethylation. As BioLCCC could only take in one gradient slope as the input, all peptides with observed RT > 48 min were not inputted into the prediction method.

ELUDE [34] was downloaded from the percolator releases page https://github.com/percolator/percolator/releases, version 3.02.0, Build Date 2018-02-02. The data were split into two, equal sets with distinct peptide sequences to form the training and test sets. The `elude` program was run with the `––no-in-source` and `––test-rt` flags. Predicted RTs from ELUDE were obtained from the testing set only, and training set RTs were not used in further analyses.

For iRT [43], the same raw files used for the previous sets were searched with the Pulsar search engine [59], with iRT alignment turned on and filtering at 1% FDR. From the Pulsar search results, only peptide sequences in common with the previous set searched in MaxQuant were selected. Predicted RT was taken from the “PP.RTPredicted” column and plotted against the empirical RT column “PP.EmpiricalRT”. Empirical RTs were not compared between those derived from MaxQuant and those derived from Pulsar.

MaxQuant match-between-runs [7, 8] was run by turning the respective option on when searching over the set of 46 experiments, and given the options of 0.7 min match time tolerance and a 20 min match time window. The “Calibrated retention time” column was used as the predicted RT, and these predicted RTs were related to observed RTs with a linear model for each experiment run.

For DART-ID, predicted RTs are the same as the mean of the inferred RT distribution, and no linear model was constructed to relate the predicted RTs to the observed RTs.

### Comparison to linear alignment model

To compare the performance of the two-piece linear model for RT alignment against a simple linear model, we ran both alignments separately on the same dataset as described in the RT alignment comparison section. For Fig. S2a, we used one experiment – 180324S_QC_SQC69A – as an example to illustrate the qualitative differences between the two models. Panels b and c used all experiments from the set to give a more quantitative comparison.

### Confidence update

We update the confidence for PSM *i* in experiment *k* according to Bayes’ theorem. Let *δ*_*ik*_ = 1 denote that PSM *i* in experiment *k* is assigned to the correct sequence (true positive), *δ*_*ik*_ = 0 denotes that the PSM is assigned to the incorrect sequence (a false positive), and as above, *ρ*_*ik*_ is an observed RT assigned to peptide *i*. At a high level, the probability that the peptide assignment is a true positive is

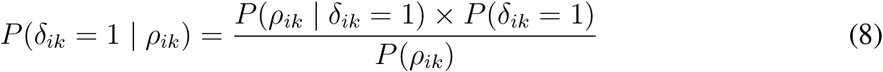

Each term is described in more detail below:

*δ*_*ik*_: An indicator for whether or not the peptide sequence assignment, *i* in experiment *k* is correct (i.e. a true or false positive).
*P*(*δ*_*ik*_ = 1|*ρ*_*ik*_): The posterior probability that the PSM is assigned to the right sequence, given the observed RT, *ρ*_*ik*_.
*P*(*ρ*_*ik*_ | *δ*_*ik*_ = 1): The RT density for peptide *i* in experiment *k* given the assignment is correct (true positive). Conditional on the alignment parameters, the true positive RT density *f*_*ik*_(*ρ*_*ik*_ | *μ*_*ik*_, *σ*_*ik*_) is 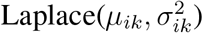. In our implementation, we incorporate uncertainty in the estimation of the alignment parameters with a parametric bootstrap, explained in more detail below and in Fig. S8.
*P*(*ρ*_*ik*_ | *δ*_*ik*_ = 0): The RT density given the assignment is incorrect (false positive). We assume that a false positive match is assigned to a peptide at random and thus take 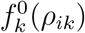 to be a broad distribution reflecting variation in *all* RTs in experiment *k*. We model this distribution as Normal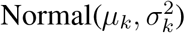, where *μ*_*k*_ is approximately the average of all RTs in the experiment and 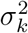 is the variance in RTs.
*P*(*δ*_*ik*_ = 1): The prior probability that the PSM’s assigned sequence is correct, i.e. one minus the posterior error probability (PEP) provided by MaxQuant, 1 − *λ*_*ik*_.
*P*(*ρ*_*ik*_): The marginal likelihood for observing the RT assigned to peptide *i* in experiment *k*. By the law of total probability, this is simply the mixture density from Eq. 5.

The confidence update depends on the global alignment parameters. Let *θ* consist of the global alignment parameters and reference RTs, i.e. *β*_0*k*_, *β*_1*k*_, *σ*_*ik*_ and *μ*_*i*_. If *θ* were known, then the Bayesian update could be computed in a straightforward manner as described above. In practice the alignment parameters are not known and thus must be estimated using the full set of observed RTs across all experiments, ****ρ****. The PSM confidence update can be expressed unconditional on *θ*, by integrating over the uncertainty in the estimates of the alignment parameters:

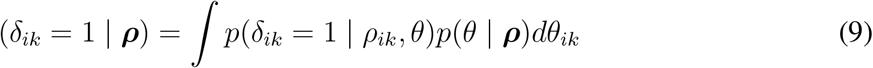

Although we can estimate this posterior distribution using Markov Chain Monte Carlo (MCMC), it is prohibitively slow given the large number of peptides and experiments that we analyze. As such, we estimate maximum a posteriori (MAP) estimates for the reference RTs *μ*_*i*_, alignment parameters *β*_0*k*_, *β*_1*k*_, and RT standard deviation *σ*_*ik*_ using an optimization routine implemented in STAN [60].

If computation time is not a concern, it is straightforward to generate posterior samples in our model by running MCMC sampling in STAN, instead of MAP optimization. This approach is computationally efficient but is limited in that parameter uncertainty quantification is not automatic.

To address this challenge, we incorporate estimation uncertainty using a computationally efficient procedure based on the parametric bootstrap. Note that uncertainty about the alignment parameters *β*_0*k*_ and *β*_1*k*_ is small since they are inferred using thousands of RT observations per experiment. By contrast, the reference RTs, *μ*_*i*_, have much higher uncertainty since we observe *at most* one RT associated with peptide *i* in each experiment (usually far fewer). As such, we choose to ignore uncertainty in the alignment parameters and focus on incorporating uncertainty in estimates of *μ*_*i*_.

Let 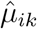 and 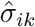 denote the MAP estimates of the location and scale parameters for the RT densities. To approximate the posterior uncertainty in the estimates of *μi*, we use the parametric bootstrap. First, we sample 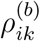 from 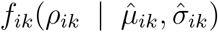 with probability 1 − *λ*_*ik*_ and 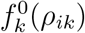 with probability *λ*_*ik*_. We then map 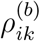 back to the reference space using the inferred alignment parameters as 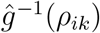 and compute a bootstrap replicate of the reference RT associated with peptide *i* as the median (across experiments) of the resampled RTs: 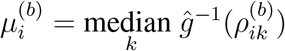, as the maximum likelihood estimate of the location parameter of a Laplace distribution is the median of independent observations. For each peptide we repeat this process *B* times to get several bootstrap replicates of the reference RT for each peptide. We use the bootstrap replicates to incorporate the uncertainty of the reference RTs into the Bayesian update of the PSM confidence. Specifically, we approximate the confidence update in Equation 9 as

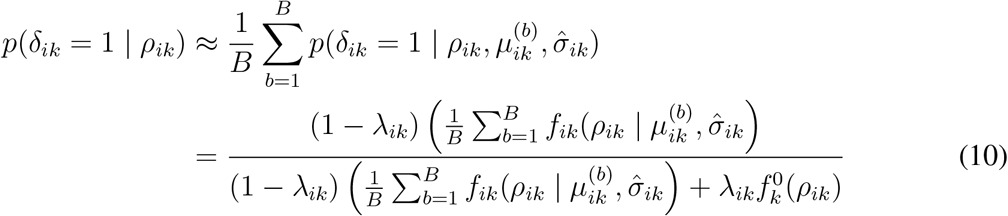

This process is depicted in Fig. S8.

In addition to updating the PEPs for each PSM, DART-ID also recalculates the set-wide false discovery rate (FDR, q-value). This is done by first sorting the PEPs and then assigning the q-value to be the cumulative sum of PEPs at that index, divided by the index itself, to give the fractional expected number of false positives at that index (i.e., the mean PEP) [55].

### TMT reporter ion intensity normalization

Reporter ion (RI) intensities were obtained by selecting the tandem-mass-tag (TMT) 11-plex labels in MaxQuant, for both attachment possibilities of lysine and the peptide N-terminus, and with a mass tolerance of 0.003 Da. Data from different experiments and searches are all combined into one matrix, where the rows are observations (PSMs) and the 10 columns are the 10 TMT channels. Observations are filtered at a confidence threshold, normally 1% FDR, and observations with missing data are thrown out.

Before normalization, empty channels 127N, 128C, and 131C are removed from the matrix. Each column of the matrix is divided by the median of that column, to correct for the total amount of protein in each channel, pipetting error, and any biases between the respective TMT tags. Then, each row of the matrix is divided by the median of that row, to obtain the relative enrichment between the samples in the different TMT channels. In our data the relative enrichment was between the two cell types present in our SCoPE-MS sets, T-cells (Jurkat cell line) and monocytes (U-937 cell lines).

Assuming that the relative RI intensities of PSMs are representative of their parent peptide, the peptide intensity can be estimated as the median of the RI intensities of its constituent PSMs. Similarly, if protein levels are assumed to correspond to the levels of its constituent peptides, then protein intensity can be estimated as the median of the intensities of its constituent peptides. The previous steps of RI normalization makes all peptide and protein-level quantitation relative between the conditions in each channel.

### Principal component analysis

For the principal component analysis as shown in Fig. 8a, data was filtered and normalized in the same manner as discussed previously. Additional experiments were manually removed from the set due to different experimental designs or poorer overall coverage that would have required additional imputation on that experiment’s inclusion.

PSMs were separated into two sets, as described in Fig. 7a: *Spectra* and *DART-ID*. PSMs in the *DART-ID* set belonging to any parent protein in the *Spectra* set were filtered out, so that the two PSM sets contained no shared proteins. Additionally, proteins that were not observed in at least 95% of the selected experiments were removed in order to reduce the amount of imputation required.

Normalized TMT quantification data was first collapsed from PSM-level to peptide-level by averaging (mean) PSM measurements for the same peptide. This process was repeated to estimate protein-level quantitation from peptide-level quantitation. This data, from both sets, was then reshaped into an expression matrix, with proteins on the rows and “single cells” (TMT channel-experiment pairs) on the columns. As described earlier in the Results section, these samples are not actual single cells but are instead comprised of cell lysate at the expected abundance of a single cell; see Table 1.

Missing values in this expression matrix were imputed with the k-nearest-neighbors (kNN) algorithm, with Euclidean distance as the similarity measure and *k* set to 5. A similarity matrix was then derived from this expression matrix by correlating (Pearson correlation) the matrix with itself. Singular value decomposition (SVD) was then performed on the similarity matrix to obtain the principal component loadings. These loadings are the left singular vectors (the columns of U of SVD: *UDV*^*T*^). Each circle was then colored based on the type of the corresponding cell from annotations of the experimental designs.

### Protein inference

Our raw data was searched with both the PSM and protein FDR threshold set, in the search engine, to 100% to include as many PSMs as possible. Therefore, once PSM confidences were updated with RT evidence, we needed to propagate those new confidences to the protein level in order to avoid any spurious protein identifications from degenerate peptide sequences [61]. This is especially pertinent as many of the new DART-ID PSMs support proteins with no other confidently identified peptides, Fig. S9. Ideally we would run our updated PSMs back through our original search engine pipeline (MaxQuant/Andromeda) [7, 21], but that is currently not possible due to technical restrictions.

Any interpretation of the DART-ID data on the protein-level was first run through the Fido protein inference algorithm [62], which gives the probability of the presence of a protein in a sample given the pool of observed peptides and the probabilities of their constituent PSMs. The Python port of Fido was downloaded from https://noble.gs.washington.edu/proj/fido and modified to be compatible with Python 3. The code was directly interfaced into DART-ID and is available to run as a user option.

For the data in this paper, protein-level analyses first had their proteins filtered at 1% FDR, where the FDR was derived from the probabilities given to each protein by the Fido algorithm. We ran Fido with the default parameters gamma: 0.5, alpha: 0.1, beta: 0.01, connected protein threshold: 14, protein grouping and using all PSMs set to false, and pruning low scores set to true.

### Application to other datasets

In Fig. 4 we evaluated DART-ID on two other third-party, publicly available datasets: iPRG 2015 [56] (MassIVE ID: MSV000079843), 12 label-free runs of yeast lysate, and TKO 2018 [57] (ProteomeXchange ID: PXD011654), 40 TMT-labelled runs of yeast lysate. Raw files were searched in MaxQuant 1.6.3.4, against a UniProt yeast database (6721 entries, 2018/05/01). The iPRG 2015 dataset was searched with cysteine carbamidomethylation (+57.02146 Da) as a fixed modification and methionine oxidation (+15.99492 Da), protein N-terminal acetylation (+42.01056 Da), and asparagine/aspartate deamidation (+0.98401 Da) as variable modifications. The TKO 2018 dataset was searched with TMT11-plex on lysine/n-terminus, cysteine carbamidomethylation (+57.02146 Da) as a fixed modification and methionine oxidation (+15.99492 Da) as a variable modification. Both searches were done with Trypsin specificity, and PSM/protein confidence thresholds were set at 1 (100%) to obtain as many PSMs as possible. Searched data, configuration files, and DART-ID analysis results are available online.

### Implementation

The DART-ID pipeline is roughly divided into three parts. First, input data from search engine output files are converted to a common format, and PSMs unsuitable for alignment are marked for removal. Second, we estimate the alignment parameters and reference RTs using an by finding the maximum of the posterior distribution (Equation 4). Initial values for the algorithm are are generated by running a simple estimation of reference RTs and linear regression parameters for *f*_*ik*_ for each experiment. Third, inferred alignment parameters and reference RTs are used to update the confidence for the PEP of a PSM.

The model was implemented using the STAN modeling language [60]. All densities were represented on the log scale. STAN was interfaced into an R script with `rstan`. STAN was used with its `optimizing` function, which gave maximum a posteriori (MAP) estimates of the parameters, as opposed to sampling from the full posterior. R was further used for data filtering, PEP updating, model adjustment, and figure creation. The code is also ported to Python3 and `pystan`, and is available as a `pip` package `dart_id` that can be run from the command-line. `DART-ID` is run with a configuration file that specifies inputs and options. All model definitions and related parameters such as distributions are defined in a modular fashion, which supports the addition of other models or fits.

Code for analysis and figure generation is available at: github.com/SlavovLab/DART-ID_2018. The `python` program for DART-ID, as well as instructions for usage and examples, are available on GitHub as a separate repository: https://github.com/SlavovLab/DART-ID. All raw files, searched data, configuration files, and analyzed data are publicly available and deposited on Mas-sIVE (ID: MSV000083149) and ProteomeXchange (ID: PXD011748).

## Supporting Information

### S1 File. DART-ID Post-run Report

A optional HTML report generated by the `dart_id` Python script. The report gives a summary of the alignment for each experiment, as well as a broad overview of the performance of the run as a whole, by showing aggregate increases in PSMs at a chosen confidence threshold. (ZIP)

### S1 Table. SCoPE-MS and mPOP Experimental Designs

An excel spreadsheet of the experimental designs of all raw files. Included are parameters for the liquid chromatography and parameters for the mass spectrometer. Also specified is the TMT channel layout for each experiment, with labels for J (T-cells, Jurkat cell line), U (monocytes, U-937 cell line), and H (human embryonic kidney cells, HEK-293 cell line). (XLSX)

### S2 Table. Mappings of raw files to figures

An excel spreadsheet providing a map that relates figures/analyses to raw files listed in Table. S1. TRUE denotes that the figure/analysis used that raw file, where FALSE denotes that it did not. (XLSX)

### S1 Figure. Mixture model incorporates spectral confidence to estimate likelihood of observing RTs

In the global alignment process, the likelihood of the alignment function and the reference RT is estimated from a mixture model, which combines the two possibilities of whether the peptide is assigned the correct or incorrect peptide sequence. These two distributions are then weighted by the error probability (PEP). This is similar to the update process, which updates the error probability and incorporates the previous error probability, as well as the two conditional probability distributions. (PDF)

### S2 Figure. Comparison of linear and segmented fits for reference RTs in experiments

(**a**) The reference RT of PSMs compared with their observed RTs, and the model plotted in green line (linear fit), or green and red lines (segmented fit, representing the two segments). For the segmented fit, the inflection point is marked with the dotted blue line. Both fits were specified separately and run separately with the same input data. (**b**) Empirical cumulative density function (ECDF) of the residual RTs for both fits. The residual RT is defined as the Observed RT − Inferred RT, where the inferred RT is the reference RT aligned to that particular experiment via. the model function – linear or segmented. (**c**) Model-fitted standard deviations, *σ*_*k*_, for each PSM as estimated by both linear and segmented fits. Points below the 45° line indicate a lower modeled RT standard deviation for the segmented fit, and vice versa. Clusters of points correspond to PSMs belonging to a particular experiment, as the PSM-specific variance of *σ*_*ik*_ is mostly reliant on the experiment in which the PSM is observed. (PDF)

### S3 Figure. Accuracy of RT inferences varies with time and between experiments

(**a**) Residual RT (observed RT –aligned RT) binned by RT for 60 min LC-MS runs. The gradient run is 5 – 35%B from 0 – 48 min, with a wash step of 35 – 100%B from 48 – 60 min. (**b**) Residual RT varying between different experiments, all 60 min LC-MS/MS runs. (PDF)

### S4 Figure. Distribution choice for inferred RT distribution and null RT distribution

(**a**) Empirical distribution of all residual RTs, i.e., Observed RT − Predicted RT, and (**b**) all RTs. Red lines denote the distributions parametrized from the data. (PDF)

### S5 Figure. Bayesian updates of PSM confidence using RTs estimated by different methods

(**a**) 2D density distributions of posterior error probabilities (PEP) derived from spectra alone (Spectral PEP) compared to the PEP after incorporating RT evidence. The RT estimates are the same as the ones shown in Fig. 2. (**b**) Comparison of updated PEP derived from DART-ID and MaxQuant RT estimates. (**c**) Increase in confident PSMs at set confidence threshold using updated PEPs. (**d**) Validation of upgraded PSMs with quantification variance within proteins. (PDF)

### S6 Figure

Distributions of Spectral PEPs and DART-ID PEPs. The bimodality of the DART-ID distribution suggests that DART-ID’s use of RTs helps cleanly separate correct from incorrect PSMs.(PDF)

### S7 Figure. Consistency of quantification between Spectra and DART-ID PSM sets

The fold change in normalized RI intensity (T-cell/monocyte), from common proteins between the *Spectra* and *DART-ID* PSM sets. We included all proteins – not just those that are significantly (< 1% FDR) differentially abundant. (PDF)

### S8 Figure. Deriving conditional probability of RT given a correct match

(**a**) The conditional probability distribution of RT given a correct peptide sequence assignment in-corporates evidence about that peptide sequence across many different experiments. “Aligned RT” is the RT after applying the alignment function, and “Std” is inferred RT standard deviation for the peptide in the given experiment. (**b**) For each RT observation for a sequence in an experiment, we infer two distributions: one corresponding to RT density given a correct PSM and the other to an incorrect PSM match. These densities are weighted by the 1-PEP and the PEP respectively and summed to produce the marginal RT distribution. (**c**) The marginal RT distribution is then used to sample B bootstrap replicates of of the observed RTs. Each bootstrapped RT is then used to construct a bootstrapped reference RT for a given sequence. The reference RT is the median of the resampled RTs (in the aligned space). (**d**) The *B* bootstrap samples of *μ*_*i*_ are used to build distributions where the variance is determined by the model-derived variance of the peptide in an experiment. (**e**) The combination of the distributions in panel (d) forms a posterior predictive distribution for the observed RT, given that the peptide sequence assignment is correct. (PDF)

### S9 Figure. Distribution of peptides quantified per protein

(**a**) Quantified PSMs per protein, including peptide sequences quantified across multiple experiments, and (**b**) peptide sequences quantified per protein. “Spectra” indicates proteins from PSMs identified below 1% FDR. “DART-ID new proteins” indicates PSMs boosted to below 1% FDR, that have different protein assignments from “Spectra”, i.e., this set of proteins and the “Spectra” set of proteins is disjoint. “DART-ID all proteins” contains all PSMs with updated DART-ID FDR < 1% FDR regardless of protein assignment. All PSMs are filtered at < 1% FDR at the protein level.(PDF)

## Acknowledgments

We thank H. Specht, T. Chen, and members of the Slavov laboratory for discussions and constructive comments. We also thank L. Reiter and L. Verbeke for access to the iRT method within the Biognosys Pulsar software. This work was funded by the National Institute of Health to N.S. under project number 1DP2GM123497-01.

## Corresponding author

Correspondence and requests for materials should be addressed to N.S.

## Author Contributions

**Conceptualization**: A.F. and N.S.

**Data Curation**: A.C.

**Formal Analysis**: A.F. and A.C.

**Funding Acquisition**: N.S.

**Investigation**: A.C., A.F. and N.S.

**Methodology**: A.F. and N.S.

**Project Administration**: N.S.

**Resources**: N.S.

**Software**: A.C. and A.F.

**Supervision**: N.S.

**Validation**: A.C.

**Visualization**: A.C.

**Writing –Original Draft Preparation**: A.C.

**Writing –Review & Editing**: A.C., A.F., and N.S.

## Supporting Information Figures

**Figure S1.**
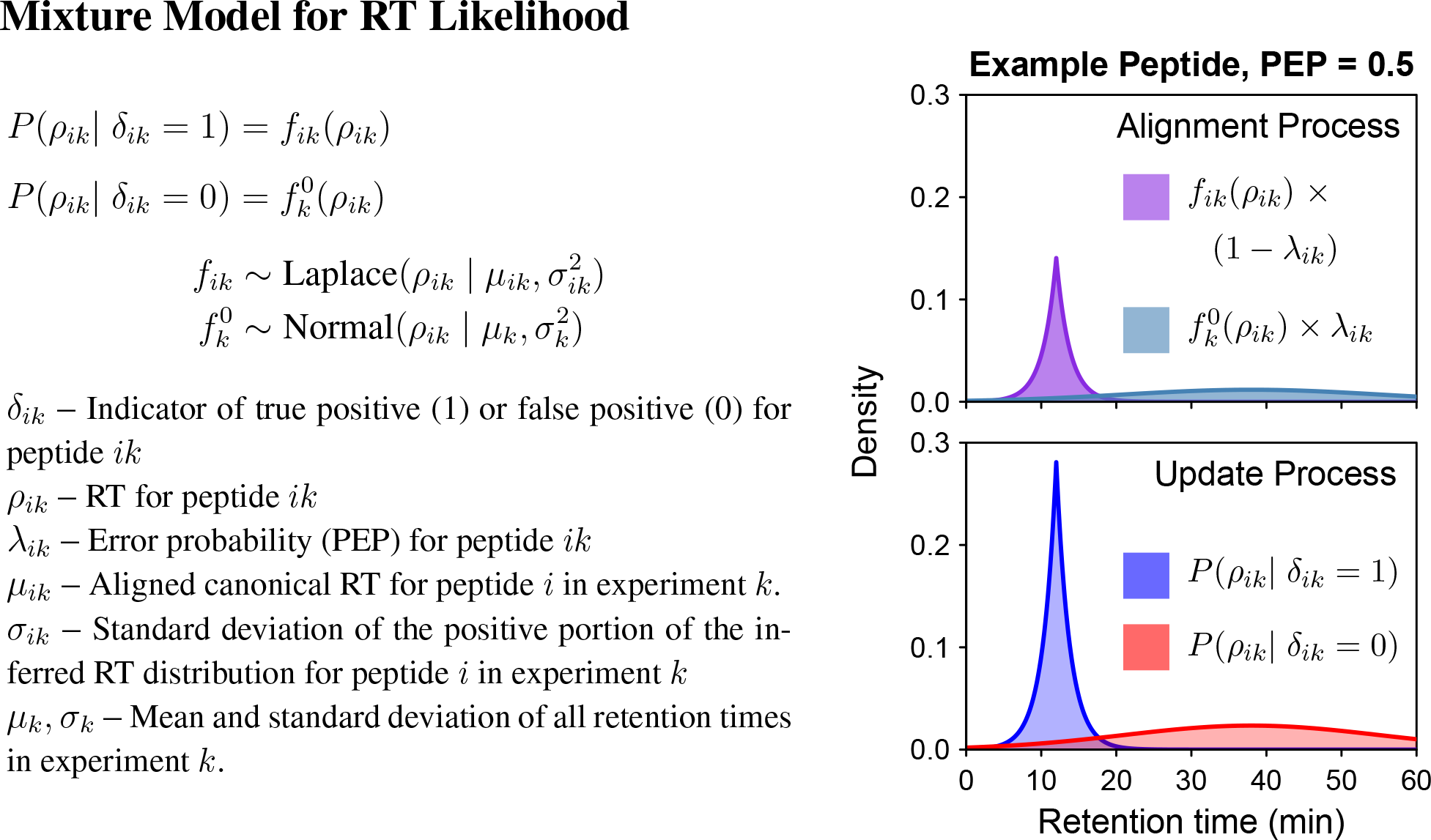
Mixture model incorporates spectral confidence to estimate likelihood of observing RTs. In the global alignment process, the likelihood of the alignment function and the reference RT is estimated from a mixture model, which combines the two possibilities of whether the peptide is assigned the correct or incorrect peptide sequence. These two distributions are then weighted by the error probability (PEP). This is similar to the update process, which updates the error probability and incorporates the previous error probability, as well as the two conditional probability distributions.

**Figure S2.**
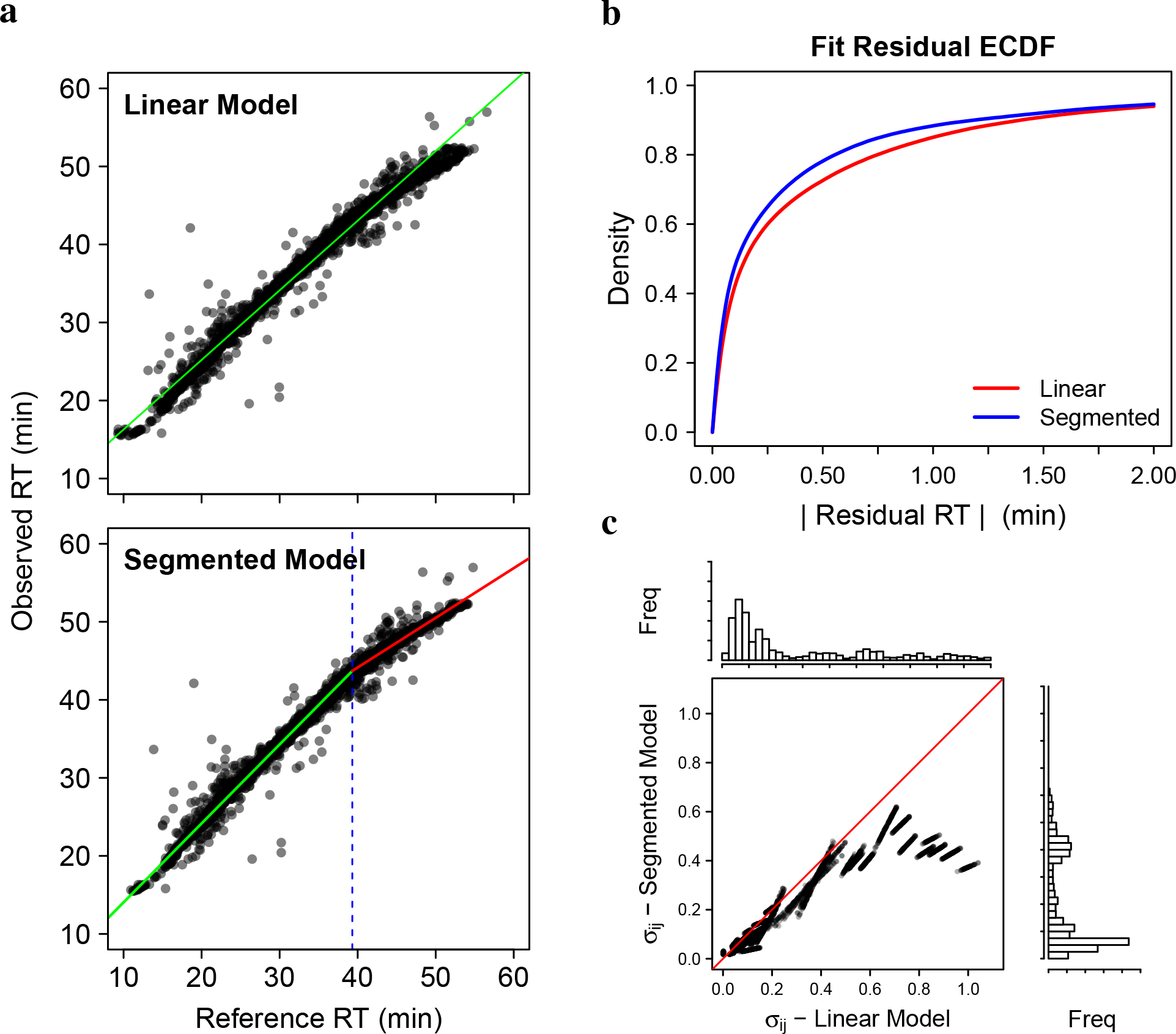
Comparison of linear and segmented fits for reference RTs in experiments. (**a**) The reference RT of PSMs compared with their observed RTs, and the model plotted in green line (linear fit), or green and red lines (segmented fit, representing the two segments). For the segmented fit, the inflection point is marked with the dotted blue line. Both fits were specified separately and run separately with the same input data. (**b**) Empirical cumulative density function (ECDF) of the residual RTs for both fits. The residual RT is defined as the Observed RT Inferred RT, where the inferred RT is the reference RT aligned to that particular experiment via. the model function – linear or segmented. (**c**) Model-fitted standard deviations, *σ*_*ik*_, for each PSM as estimated by both linear and segmented fits. Points below the 45° line indicate a lower modeled RT standard deviation for the segmented fit, and vice versa. Clusters of points correspond to PSMs belonging to a particular experiment, as the PSM-specific variance of *σ*_*ik*_ is mostly reliant on the experiment in which the PSM is observed.

**Figure S3.**
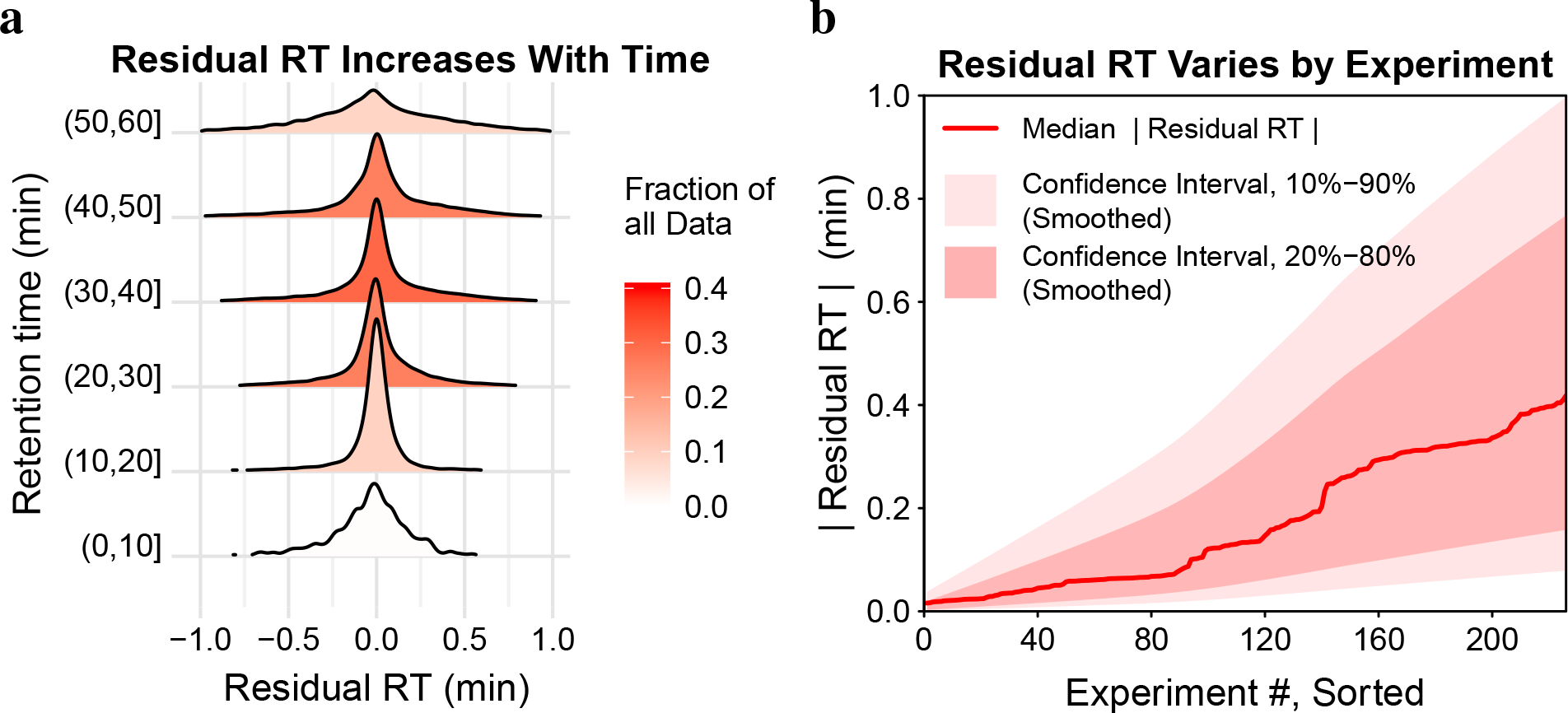
Accuracy of RT inferences varies with time and between experiments. (**a**) Residual RT (observed RT – aligned RT) binned by RT for 60 min LC-MS runs. The gradient run is 5 – 35%B from 0 – 48 min, with a wash step of 35 – 100%B from 48 – 60 min. (**b**) Residual RT varying between different experiments, all 60 min LC-MS/MS runs.

**Figure S4.**
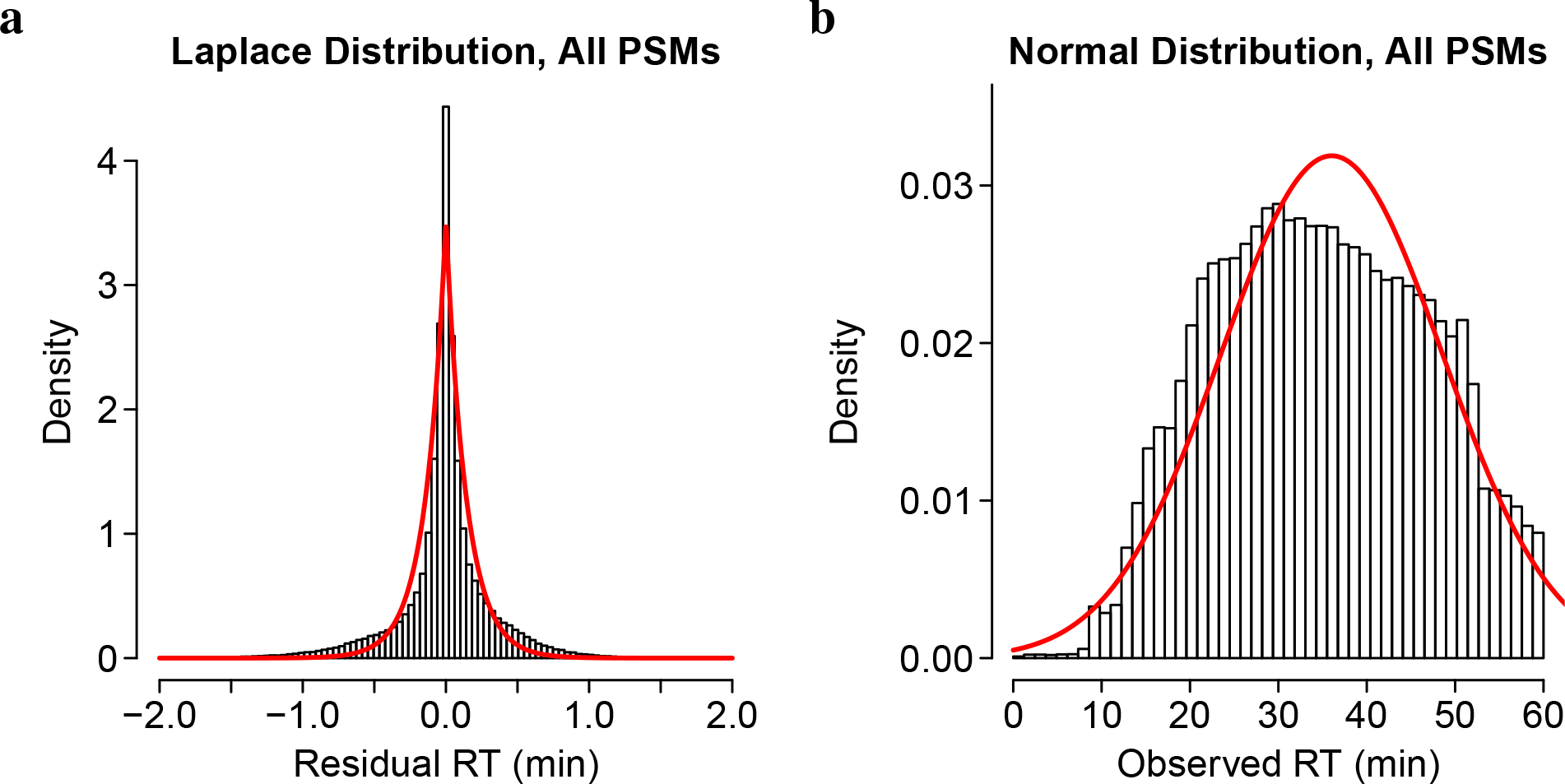
Distribution choice for inferred RT distribution and null RT distribution. (**a**) Empirical distribution of all residual RTs, i.e., Observed RT — Predicted RT, and (**b**) all RTs. Red lines denote the distributions parametrized from the data.

**Figure S5.**
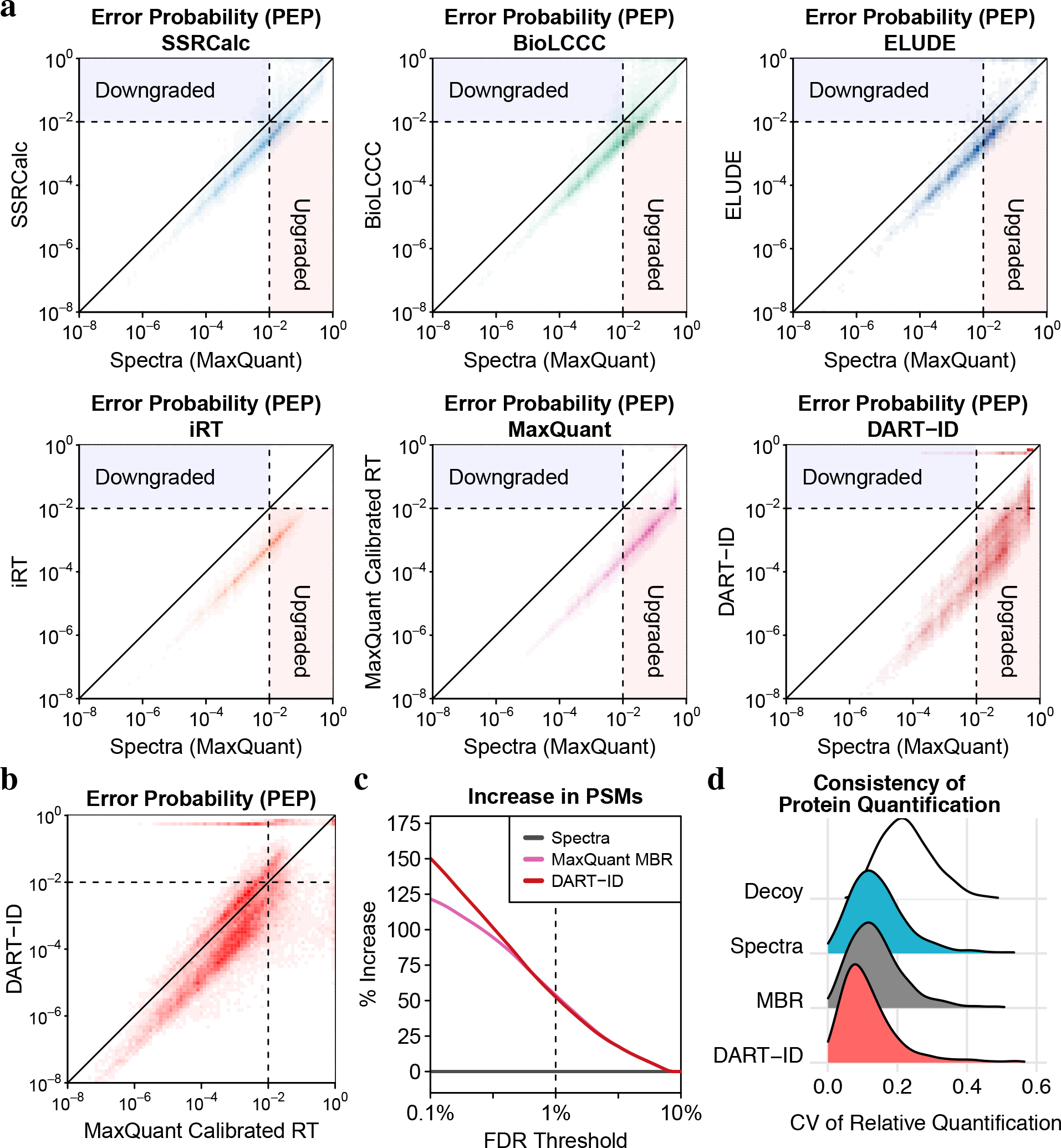
Bayesian updates of PSM confidence using RTs estimated by different methods. (**a**) 2D density distributions of posterior error probabilities (PEP) derived from spectra alone (Spectral PEP) compared to the PEP after incorporating RT evidence. The RT estimates are the same as the ones shown in Fig. 2. (**b**) Comparison of updated PEP derived from DART-ID and MaxQuant RT estimates. (**c**) Increase in confident PSMs at set confidence threshold using updated PEPs. (**d**) Validation of upgraded PSMs with quantification variance within proteins.

**Figure S6.**
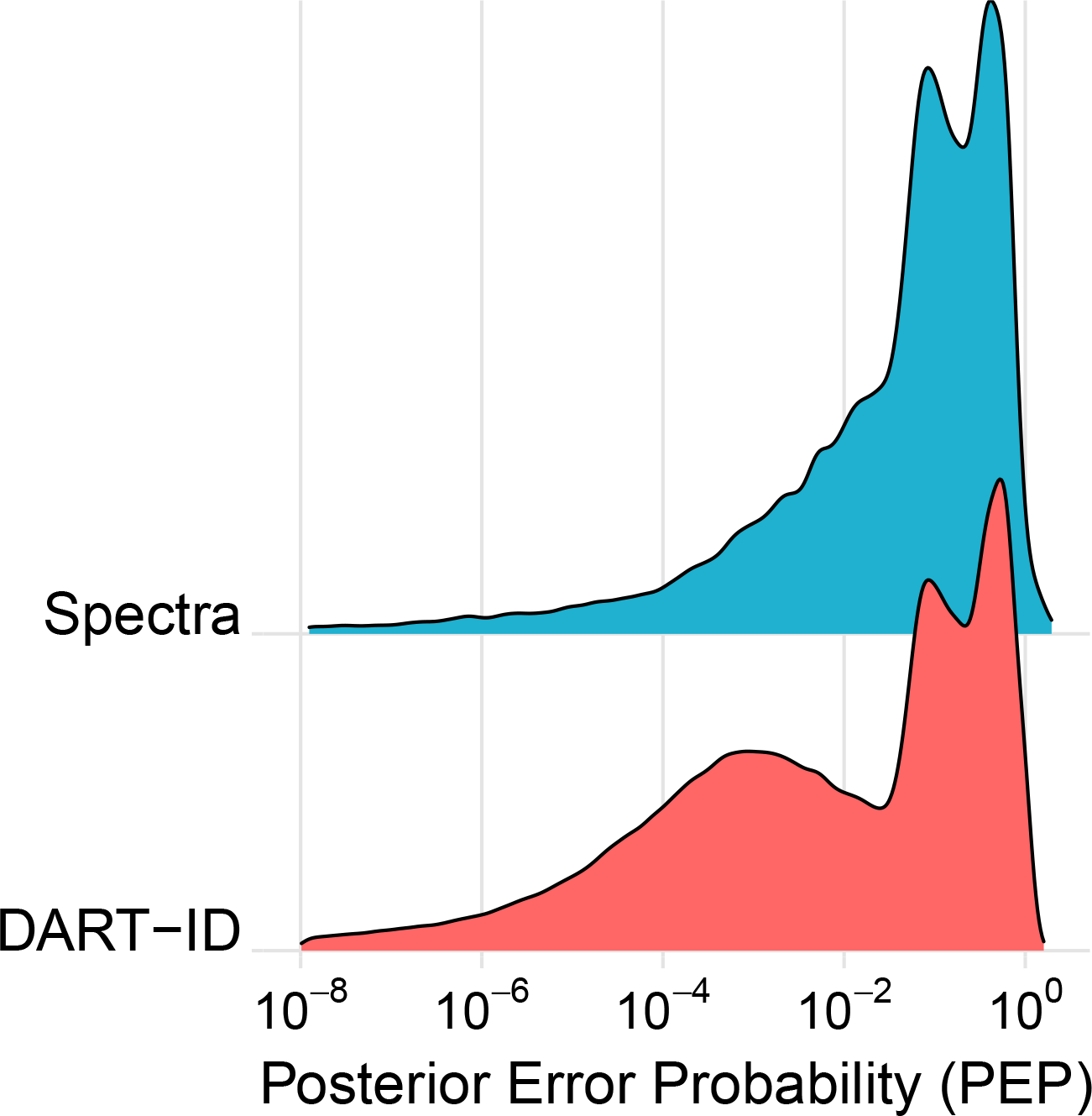
Distributions of Spectral PEPs and DART-ID PEPs. The bimodality of the DART-ID distribution suggests that DART-ID’s use of RTs helps cleanly separate correct from incorrect PSMs.

**Figure S7.**
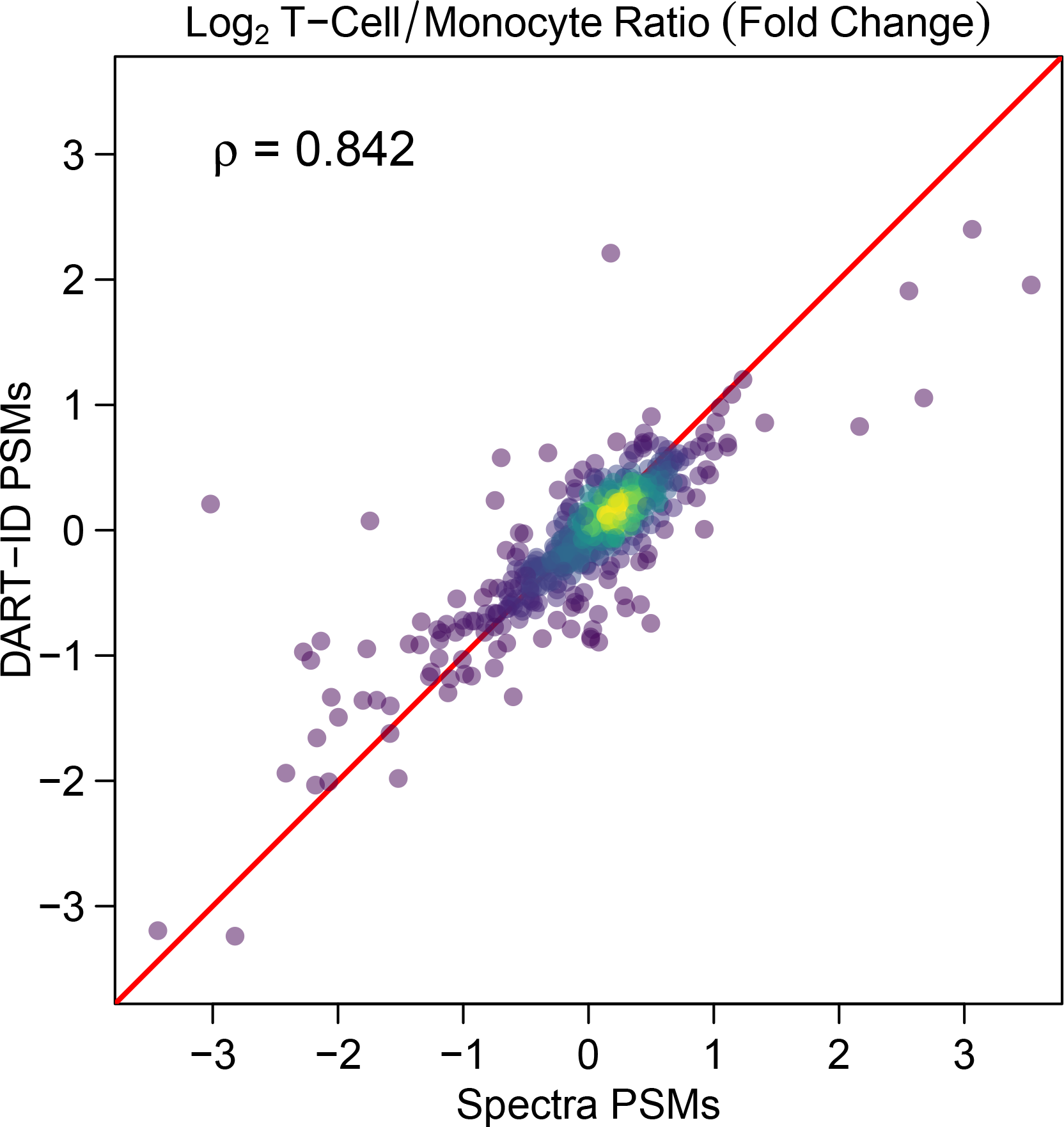
Consistency of quantification between Spectra and DART-ID PSM sets. The fold change in normalized RI intensity (T-cell/monocyte), from common proteins between the *Spectra* and *DART-ID* PSM sets. We included all proteins – not just those that are significantly (< 1% FDR) differentially abundant.

**Figure S8.**
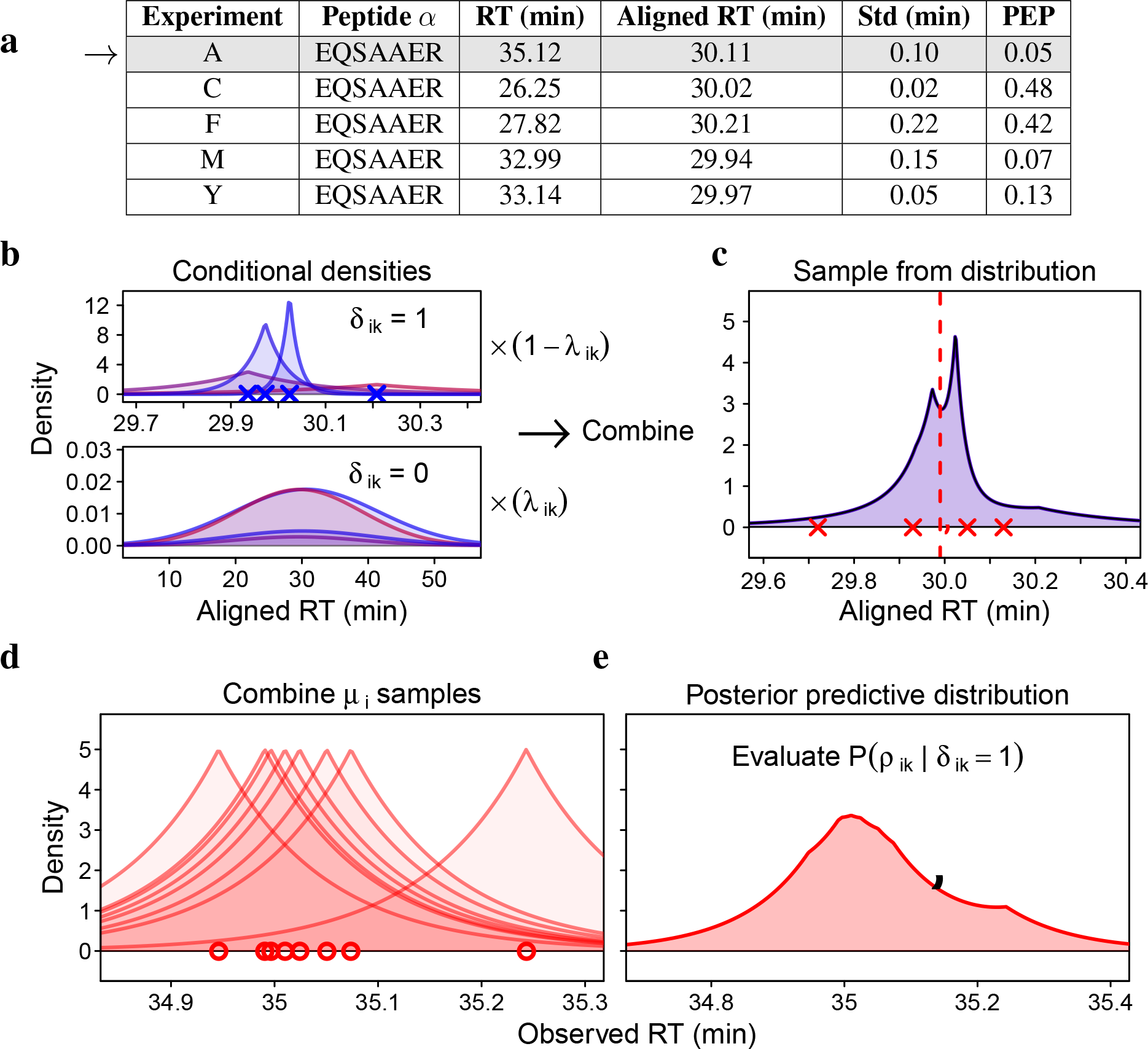
Deriving conditional probability of RT given a correct match. (**a**) The conditional probability distribution of RT given a correct peptide sequence assignment incorporates evidence about that peptide sequence across many different experiments. “Aligned RT” is the RT after applying the alignment function, and “Std” is inferred RT standard deviation for the peptide in the given experiment. (**b**) For each RT observation for a sequence in an experiment, we infer two distributions: one corresponding to RT density given a correct PSM and the other to an incorrect PSM match. These densities are weighted by the 1-PEP and the PEP respectively and summed to produce the marginal RT distribution. (**c**) The marginal RT distribution is then used to sample B bootstrap replicates of of the observed RTs. Each bootstrapped RT is then used to construct a bootstrapped reference RT for a given sequence. The reference RT is the median of the resampled RTs (in the aligned space). (**d**) The *B* bootstrap samples of *μ*_*i*_ are used to build distributions where the variance is determined by the model-derived variance of the peptide in an experiment. (**e**) The combination of the distributions in panel (d) forms a posterior predictive distribution for the observed RT, given that the peptide sequence assignment is correct.

**Figure S9.**
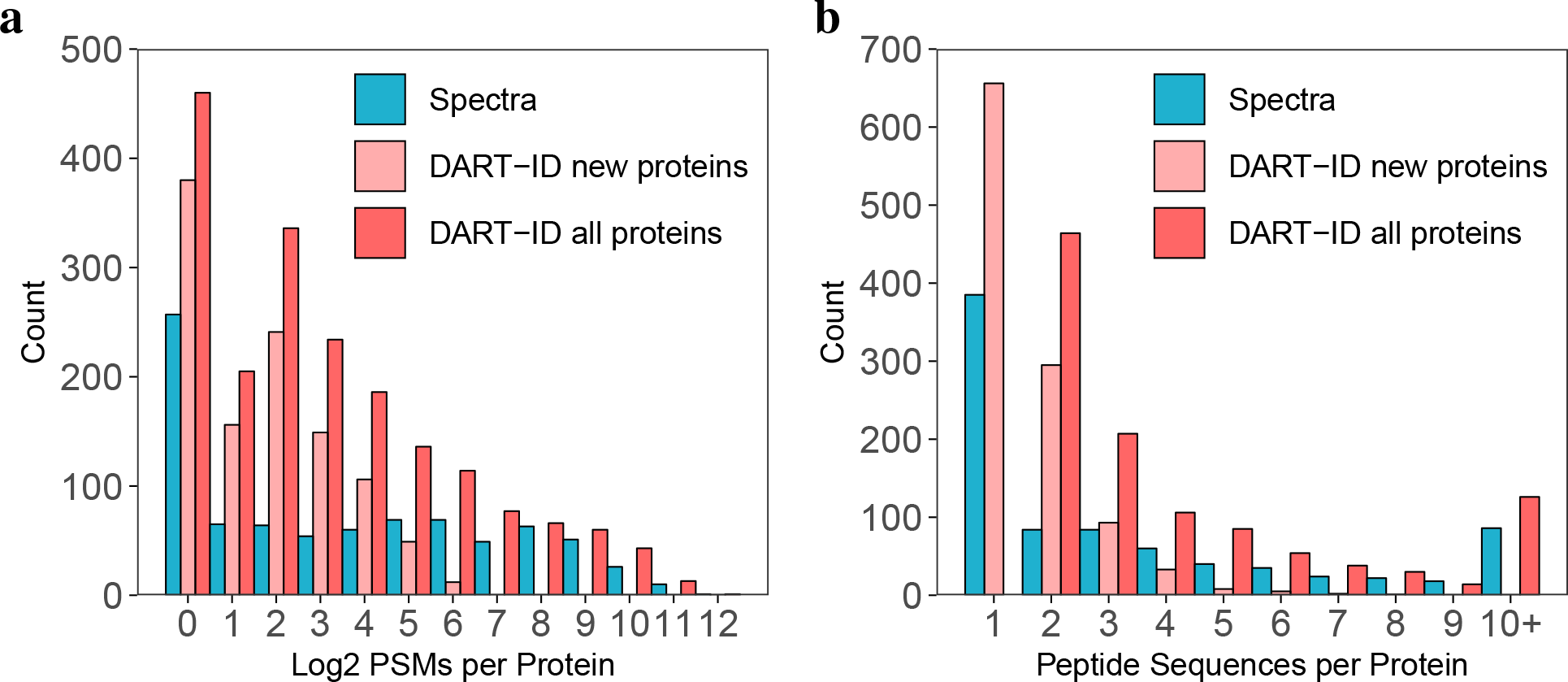
Distribution of peptides quantified per protein. Quantified PSMs per protein, including peptide sequences quantified across multiple experiments, and (**b**) peptide sequences quantified per protein. “Spectra” indicates proteins from PSMs identified below 1% FDR. “DART-ID new proteins” indicates PSMs boosted to below 1% FDR, that have different protein assignments from “Spectra”, i.e., this set of proteins and the “Spectra” set of proteins is disjoint. “DART-ID all proteins” contains all PSMs with updated DART-ID FDR < 1% FDR regardless of protein assignment. All PSMs are filtered at < 1% FDR at the protein level.

## References

1. Budnik, B., Levy, E., Harmange, G. & Slavov, N. SCoPE-MS: mass-spectrometry of single mammalian cells quantifies proteome heterogeneity during cell differentiation. Genome Biology 19, 161 (2018).

2. Specht, H. et al. Automated sample preparation for high-throughput single-cell proteomics. bioRxiv. doi:10.1101/399774 (2018).

3. Levy, E. & Slavov, N. Single cell protein analysis for systems biology. Essays In Biochemistry 62. doi:10.1042/EBC20180014 (2018).

4. Specht, H. & Slavov, N. Transformative opportunities for single-cell proteomics. Journal of Proteome Research 17, 2563–2916 (8 June 2018).

5. MacLean, B. et al. Skyline: an open source document editor for creating and analyzing targeted proteomics experiments. Bioinformatics 26, 966–968 (2010).

6. Argentini, A. et al. moFF: a robust and automated approach to extract peptide ion intensities. en. Nature Methods 13, 964–966. ISSN: 1548-7091, 1548-7105 (Dec. 2016).

7. Cox, J. & Mann, M. MaxQuant enables high peptide identification rates, individualized p.p.b.-range mass accuracies and proteome-wide protein quantification. Nature Biotechnology 26, 1367–1372 (2008).

8. Tyanova, S., Temu, T. & Cox, J. The MaxQuant computational platform for mass spectrometry-based shotgun proteomics. Nature Protocols 11, 2301–2319 (2016).

9. Zhang, B., Käll, L. & Zubarev, R. A. DeMix-Q: Quantification-Centered Data Processing Workflow. en. Molecular & Cellular Proteomics 15, 1467–1478. ISSN: 1535-9476, 1535-9484 (Apr. 2016).

10. Weisser, H. & Choudhary, J. S. Targeted Feature Detection for Data-Dependent Shotgun Proteomics. en. Journal of Proteome Research 16, 2964–2974. ISSN: 1535-3893, 1535-3907 (Aug. 2017).

11. Cox, J. et al. Accurate Proteome-wide Label-free Quantification by Delayed Normalization and Maximal Peptide Ratio Extraction, Termed MaxLFQ. en. Molecular & Cellular Proteomics 13, 2513–2526. ISSN: 1535-9484 (June 2014).

12. Ong, S.-E. & Mann, M. A practical recipe for stable isotope labeling by amino acids in cell culture (SILAC). en. Nature Protocols 1, 2650–2660. ISSN: 1754-2189 (Jan. 2007).

13. Käll, L., Canterbury, J. D., Weston, J., Noble, W. S. & MacCoss, M. J. Semi-supervised learning for peptide identification from shotgun proteomics datasets. Nature Methods 4, 923–925 (2007).

14. Strittmatter, E. F. et al. Application of Peptide LC Retention Time Information in a Discriminant Function for Peptide Identification by Tandem Mass Spectrometry. Journal of Proteome Research 3, 760–769 (4 2004).

15. Klammer, A. A., Yi, X., MacCoss, M. J. & Noble, W. S. Improving Tandem Mass Spectrum Identification Using Peptide Retention Time Prediction across Diverse Chromatography Conditions. Analytical Chemistry 79.PMID: 17622186, 6111–6118 (2007).

16. Pfeifer, N., Leinenbach, A., Huber, C. G. & Kohlbacher, O. Statistical learning of peptide retention behavior in chromatographic separations: a new kernel-based approach for computational proteomics. en. BMC Bioinformatics 8, 468. ISSN: 1471-2105 (2007).

17. Pfeifer, N., Leinenbach, A., Huber, C. G. & Kohlbacher, O. Improving Peptide Identification in Proteome Analysis by a Two-Dimensional Retention Time Filtering Approach. en. Journal of Proteome Research 8, 4109–4115. ISSN: 1535-3893, 1535-3907 (Aug. 2009).

18. Li, G.-Z. et al. Database searching and accounting of multiplexed precursor and product ion spectra from the data independent analysis of simple and complex peptide mixtures. en. Proteomics 9, 1696–1719. ISSN: 16159853, 16159861 (Mar. 2009).

19. Dorfer, V., Maltsev, S., Winkler, S. & Mechtler, K. CharmeRT: Boosting peptide identifications by chimeric spectra identification and retention time prediction. Journal of Proteome Research 0.PMID: 29863353, null (2018).

20. Keller, A., Nesvizhskii, A. I., Kolker, E. & Aebersold, R. Empirical Statistical Model To Estimate the Accuracy of Peptide Identifications Made by MS/MS and Database Search. Analytical Chemistry 74, 5383–5392 (2002).

21. Cox, J. et al. Andromeda: A Peptide Search Engine Integrated into the MaxQuant Environment. en. Journal of Proteome Research 10, 1794–1805. ISSN: 1535-3893, 1535-3907 (Apr. 2011).

22. Moruz, L. & Käll, L. Peptide retention time prediction. en. Mass Spectrometry Reviews 36, 615–623. ISSN: 02777037 (Sept. 2017).

23. Krokhin, O. V. et al. Use of Peptide Retention Time Prediction for Protein Identification by off-line Reversed-Phase HPLC-MALDI MS/MS. en. Analytical Chemistry 78, 6265–6269. ISSN: 0003-2700, 1520-6882 (Sept. 2006).

24. McQueen, P. et al. Information-dependent LC-MS/MS acquisition with exclusion lists potentially generated on-the-fly: Case study using a whole cell digest of *Clostridium thermocellum*. en. PROTEOMICS 12, 1160–1169. ISSN: 16159853 (Apr. 2012).

25. Meek, J. L. Prediction of peptide retention times in high-pressure liquid chromatography on the basis of amino acid composition. en. Proceedings of the National Academy of Sciences 77, 1632–1636. ISSN: 0027-8424, 1091-6490 (Mar. 1980).

26. Guo, D., Mant, C. T., Taneja, A. K., Parker, J. & Rodges, R. S. Prediction of peptide retention times in reversed-phase high-performance liquid chromatography I. Determination of retention coefficients of amino acid residues of model synthetic peptides. Journal of Chromatography A 359, 499–518. ISSN: 0021-9673 (1986).

27. Sakamoto, Y., Kawakami, N. & Sasagawa, T. Prediction of peptide retention times. en. Journal of Chromatography A 442, 69–79. ISSN: 00219673 (Jan. 1988).

28. Krokhin, O. et al. An Improved Model for Prediction of Retention Times of Tryptic Peptides in Ion Pair Reversed-phase HPLC: Its Application to Protein Peptide Mapping by Off-Line HPLC-MALDI MS. en. Molecular & Cellular Proteomics 3, 908–919. ISSN: 1535-9476, 1535-9484 (Sept. 2004).

29. Baczek, T., Wiczling, P., Marszall, M., Heyden, Y. V. & Kaliszan, R. Prediction of Peptide Retention at Different HPLC Conditions from Multiple Linear Regression Models. en. Journal of Proteome Research 4, 555–563. ISSN: 1535-3893, 1535-3907 (Apr. 2005).

30. Krokhin, O. V. Sequence-Specific Retention Calculator. Algorithm for Peptide Retention Prediction in Ion-Pair RP-HPLC: Application to 300- and 100-Å Pore Size C18 Sorbents. Analytical Chemistry 78.PMID: 17105172, 7785–7795 (2006).

31. Gorshkov, A. V. et al. Liquid Chromatography at Critical Conditions: Comprehensive Approach to Sequence-Dependent Retention Time Prediction. en. Analytical Chemistry 78, 7770–7777. ISSN: 0003-2700, 1520-6882 (Nov. 2006).

32. Petritis, K. et al. Use of Artificial Neural Networks for the Accurate Prediction of Peptide Liquid Chromatography Elution Times in Proteome Analyses. en. Analytical Chemistry 75, 1039–1048. ISSN: 0003-2700, 1520-6882 (Mar. 2003).

33. Petritis, K. et al. Improved Peptide Elution Time Prediction for Reversed-Phase Liquid Chromatography-MS by Incorporating Peptide Sequence Information. Analytical Chemistry 78, 5026–5039 (2006).

34. Moruz, L., Tomazela, D. & Käll, L. Training, Selection, and Robust Calibration of Retention Time Models for Targeted Proteomics. Journal of Proteome Research 9.PMID: 20735070, 5209–5216 (2010).

35. Lu, W. et al. Locus-specific Retention Predictor (LsRP): A Peptide Retention Time Predictor Developed for Precision Proteomics. en. Scientific Reports 7, 43959. ISSN: 2045-2322 (Mar. 2017).

36. Palmblad, M., Ramström, M., Markides, K. E., Hãkansson, P. & Bergquist, J. Prediction of Chromatographic Retention and Protein Identification in Liquid Chromatography/Mass Spectrometry. en. Analytical Chemistry 74, 5826–5830. ISSN: 0003-2700, 1520-6882 (Nov. 2002).

37. Palmblad, M. et al. Protein identification by liquid chromatography-mass spectrometry using retention time prediction. en. Journal of Chromatography B 803, 131–135. ISSN: 15700232 (Apr. 2004).

38. Silva, J. C. et al. Quantitative Proteomic Analysis by Accurate Mass Retention Time Pairs. Analytical Chemistry 77.PMID: 15801753, 2187–2200 (2005).

39. Conrads, T. P., Anderson, G. A., Veenstra, T. D., Paša-Tolić, L. & Smith, R. D. Utility of Accurate Mass Tags for Proteome-Wide Protein Identification. en. Analytical Chemistry 72, 3349–3354. ISSN: 0003-2700, 1520-6882 (July 2000).

40. Norbeck, A. D. et al. The Utility of Accurate Mass and LC Elution Time Information in the Analysis of Complex Proteomes. en. Journal of the American Society for Mass Spectrometry 16, 1239–1249. ISSN: 1044-0305, 1879-1123 (Aug. 2005).

41. Bochet, P. et al. Fragmentation-free LC-MS can identify hundreds of proteins. en. Proteomics 11, 22–32. ISSN: 16159853 (Jan. 2011).

42. Krokhin, O. V. & Spicer, V. Peptide Retention Standards and Hydrophobicity Indexes in Reversed-Phase High-Performance Liquid Chromatography of Peptides. en. Analytical Chemistry 81, 9522–9530. ISSN: 0003-2700, 1520-6882 (Nov. 2009).

43. Escher, C. et al. Using iRT, a normalized retention time for more targeted measurement of peptides. Proteomics 12, 1111–1121. ISSN: 1615-9861 (2012).

44. Van Nederkassel, A., Daszykowski, M., Eilers, P. & Heyden, Y. V. A comparison of three algorithms for chromatograms alignment. en. Journal of Chromatography A 1118, 199–210. ISSN: 00219673 (June 2006).

45. Podwojski, K. et al. Retention time alignment algorithms for LC/MS data must consider non-linear shifts. Bioinformatics 25, 758–764 (2009).

46. Lange, E., Tautenhahn, R., Neumann, S. & Gröpl, C. Critical assessment of alignment procedures for LC-MS proteomics and metabolomics measurements. en. BMC Bioinformatics 9, 375. ISSN: 1471-2105 (2008).

47. Stanstrup, J., Neumann, S. & Vrhovšek, U. PredRet: Prediction of Retention Time by Direct Mapping between Multiple Chromatographic Systems. en. Analytical Chemistry 87, 9421–9428. ISSN: 0003-2700, 1520-6882 (Sept. 2015).

48. Fischer, B. et al. Semi-supervised LC/MS alignment for differential proteomics. en. Bioinformatics 22, e132–e140. ISSN: 1367-4803, 1460-2059 (July 2006).

49. Bernhardt, O. M. et al. Spectronaut A fast and efficient algorithm for MRM-like processing of data independent acquisition (SWATH-MS) data. en, 1 (2012).

50. Gillet, L. C. et al. Targeted Data Extraction of the MS/MS Spectra Generated by Data-independent Acquisition: A New Concept for Consistent and Accurate Proteome Analysis. en. Molecular & Cellular Proteomics 11, 17 (Jan. 2012).

51. Röst, H. L. et al. OpenSWATH enables automated, targeted analysis of data-independent acquisition MS data. en. Nature Biotechnology 32, 219–223. ISSN: 1087-0156, 1546-1696 (Mar. 2014).

52. Bruderer, R., Bernhardt, O. M., Gandhi, T. & Reiter, L. High-precision iRT prediction in the targeted analysis of data-independent acquisition and its impact on identification and quantitation. en. PROTEOMICS 16, 2246–2256. ISSN: 16159853 (Aug. 2016).

53. Malioutov, D. & Slavov, N. Convex Total Least Squares. Journal of Machine Learning Research 32, 109–117 (1 2014).

54. Elias, J. E. & Gygi, S. P. Target-decoy search strategy for increased confidence in large-scale protein identifications by mass spectrometry. en. Nature Methods 4, 207–214. ISSN: 1548-7091, 1548-7105 (Mar. 2007).

55. Käll, L., Storey, J. D., MacCoss, M. J. & Noble, W. S. Posterior Error Probabilities and False Discovery Rates: Two Sides of the Same Coin. en. Journal of Proteome Research 7, 40–44. ISSN: 1535-3893, 1535-3907 (Jan. 2008).

56. Choi, M. et al. ABRF Proteome Informatics Research Group (iPRG) 2015 Study: Detection of Differentially Abundant Proteins in Label-Free Quantitative LC–MS/MS Experiments. en. Journal of Proteome Research 16, 945–957. ISSN: 1535-3893, 1535-3907 (Feb. 2017).

57. Gygi, J. P. et al. Web-Based Search Tool for Visualizing Instrument Performance Using the Triple Knockout (TKO) Proteome Standard. en. Journal of Proteome Research. ISSN: 1535-3893, 1535-3907. doi:10.1021/acs.jproteome.8b00737 (Nov. 2018).

58. Franks, A., Airoldi, E. & Slavov, N. Post-transcriptional regulation across human tissues. PLOS Computational Biology 13, e1005535 (2017).

59. Verbeke, L., Bernhardt, O. M., Gandhi, T., Bruderer, R. & Reiter, L. Pulsar: A Search Engine Integrated into Spectronaut using Dynamic PSM Stratification. en, 1 (2017).

60. Carpenter, B. et al. Stan: A Probabilistic Programming Language 2017. doi:10.18637/jss.v076.i01.

61. Nesvizhskii, A. I., Keller, A., Kolker, E. & Aebersold, R. A Statistical Model for Identifying Proteins by Tandem Mass Spectrometry. en. Analytical Chemistry 75, 4646–4658. ISSN: 0003-2700, 1520-6882 (Sept. 2003).

62. Serang, O., MacCoss, M. J. & Noble, W. S. Efficient Marginalization to Compute Protein Posterior Probabilities from Shotgun Mass Spectrometry Data. en. Journal of Proteome Research 9, 5346–5357. ISSN: 1535-3893, 1535-3907 (Oct. 2010).

